# Evolution, Expansion and Characterization of Cannabinoid Synthase Gene Family in *Cannabis Sativa*

**DOI:** 10.1101/2022.11.18.517131

**Authors:** Keith D Allen, Anthony Torres, Kymron De Cesare, Reginald Gaudino

**Affiliations:** Front Range Biosciences –; Front Range Biosciences -; Front Range Biosciences

## Abstract

We are working toward a complete functional and genomic characterization of the cannabinoid synthase family in *Cannabis* (itself part of the larger Berberine Bridge Enzyme family). This clade, which is unique to *Cannabis*, includes four main subclades that appear to have arisen by a series of gene duplications. We have functionally characterized three cannabinoid synthases, in addition to the three already characterized, so that now all four subclades contain at least one characterized enzyme. The previously uncharacterized Clade C enzymes have relatively low activity and produce CBCA as their primary product. In addition, we report genomic characterization to better understand the structure and high level of copy number variation in this family. We report a pattern of shared regions immediately upstream of the cannabinoid synthase genes that suggests a specific sequence of breakpoints, and hence gene duplication events. We present a model of gene family expansion to account for the observed data, along with evidence that this expansion occurred under selective pressure. This work adds to our understanding of both the high level of variability in this family and the origin of THCA in type III plants lacking a functional THCAS gene.

## Introduction

Cannabinoids, defined chemically as isoprenylated resorcinol polyketides, are best known in *Cannabis sativa*, where more than a 120 have been identified, but interestingly, they are also found in a range of higher plants, liverwort, and even fungi (Hanuš et al., 2016; Gülck and Møller, 2020). *Cannabis* is one of the earliest domesticated plants, grown for fiber, seed protein, and psychoactive properties for thousands of years. Wide sequencing of diverse *Cannabis* accessions indicates that there has been strong selective pressure throughout this domestication period (Ren, et al 2021). While the total number of cannabinoids is bewilderingly large, the main constituents are Δ9-tetrahydrocannabinolic acid (THCA), which is primarily responsible for the psychotropic aspects of this plant, and cannabidiolic acid (CBDA), which while not psychotropic has a range of potential medical applications, including very promising anticonvulsant properties (Lazarini-Lopes et al., 2020). Of the less well studied cannabinoids, cannabichromenic acid (CBCA) is the most abundant, but while this compound is commonly detectable in medicinal or recreational samples, concentrations in flower material are typically less than 0.5% of dry weight. The two most abundant cannabinoids, THCA and CBDA define the chemotype of this plant with THCA dominant accessions labeled Type I, or drug-type, accessions with an approximately 1:1 ratio of THCA:CBDA defined as Type II, and CBDA dominant accessions defined as Type III, or hemp type. The Hemp Farming Act of 2018 defined hemp as containing less than 0.3% THC, creating a legal distinction between hemp and drug-type *Cannabis*, which is still classified as a schedule I narcotic. This allows legal cultivation of hemp for fiber or seed protein production, but also for CBDA production, given that CBDA is not a controlled substance, and preparations of this compound have now been approved by the FDA for treatment of epileptic seizures (Michael Felberbaum, 2018; Pauli et al., 2020). Demand for CBDA has been steadily rising as new possible uses emerge, leading to an industry growing hemp to satisfy this demand. But even with Type III hemp plants that can be shown to lack a functional THCA synthase gene, THCA is still present in these plants at a fairly constant ratio of around 24-28 to 1 CBDA:THCA, and the relationship is linear with a Pearson’s r = 0.98 (Zirpel et al., 2018; Toth et al., 2020). This makes it quite difficult to grow plants with a CBDA concentration above about 8% of dry weight without THCA concentrations rising above the 0.3% threshold, rendering the crop out of compliance (Stack et al., 2021). One of the many unanswered questions in this species is the origin of THCA in plants that lack a functional THCAS gene, with some suggestions that it could be produced as a side product of one or more of the other related synthases, in particular CBCA synthases (Onofri et al., 2015). This remains a significant unsolved problem.

The genes encoding the enzymes responsible for production of THCA, CBDA, and CBCA have all been identified and characterized (Sirikantaramas et al., 2004; Taura et al., 2007; Laverty et al., 2019). And the crystal structure of THCA synthase (THCAS) has been solved (Shoyama et al., 2012). The genes encoding CBDAS and THCAS appear to be tightly linked genetically, and initial data on the segregation of chemical phenotype was consistent with a simple Mendelian model involving codominant alleles at a single locus (de Meijer et al., 2003). As the genes were sequenced and improved assemblies were published, it became clear that these are separate genes, but tightly linked enough to behave as a single locus (van Bakel et al., 2011). It is now well established that chemotype is determined by two tightly linked genes that are in repulsion, one chromosome with an active THCAS and an inactive CBDAS, while the other chromosome has the opposite arrangement (Weiblen et al., 2015; Grassa et al., 2021). This arrangement would explain why neither of the assemblies from type II accessions (Cannbio2, GCA_016165845.1 and Jamaican Lion, GCA_003660325.2) successfully assembled intact THCAS and CBDAS genes on the same contig.

The cannabinoid synthases are encoded by single exon genes with proteins of about 545 amino acids, including a 28 amino acid N-terminal secretion signal. These enzymes have been classified as oxidocyclases belonging to the evolutionarily ancient berberine bridge enzyme family (BBE, Pfam PF08031), a family characterized by relatively low sequence conservation and an astonishing diversity of catalyzed chemical reactions (Daniel et al., 2017). These enzymes all share a common fold, and an unusual bivalently linked FAD molecule. While enzymes in this family are known to produce a diverse array of alkaloids, the vast majority of identified genes have not been characterized.

Using a phylogenetic approach, it has been shown that cannabinoid oxidocyclase genes are specific to *Cannabis*, and that this section of the BBE family arose from a single progenitor gene in an ancestral *Cannabis* plant, rather than originating from an earlier duplication event prior to *Cannabis* speciation (van Velzen and Schranz, 2021). While the origin of this gene family is beginning to be elucidated, there is still much to learn about how this family evolved in *Cannabis*, and what role human selection may have played in that evolution.

Once thought to only consist of synthases for THCA and CBDA and a variable number of pseudogenes, or “inactive THCA-like synthase” genes, this family can have as many as a dozen actively expressed and functional genes in a given accession. But a key feature of this family is that the number of genes is quite variable from accession to accession. (van Velzen and Schranz, 2021). As many as a hundred cannabinoids have been detected in *Cannabis*, but the enzymes characterized so far only account for the most abundant molecules, and a few unknowns (Hanuš et al., 2016). Given the amount of medicinal potential, it is important to understand how many of the enzymes in this family are active, and the products they produce. In this paper we characterize three more of these enzymes, substantially adding to our knowledge of product profiles produced by this family. Particularly, this work helps us understand the enzymes responsible for THCA production in CBDA-dominant hemp lines that only contain an inactive THCAS, which is of importance to hemp farmers and breeders to maintain compliance. We also describe genomic signatures suggesting a history of the expansion of this family after speciation of Cannabis.

## Methods and Materials

### Genomic DNA Isolation

Genomic DNA was isolated from *Cannabis* samples using the Qiagen DNA Easy Plant genomic DNA isolation kits (Qiagen) using manufacturer instructions for preparation of genomic DNA Extracts. Purified DNA was prepared for quantification using the Qubit HS-dsDNA System (Promega) quantified using a Qubit Fluorometer (Thermo Fisher) as per manufacturer’s instructions. Quantitated DNA was diluted to 5ng/uL final working concentration and used as normalized input for PCR reactions.

### Degenerate Primer Set for Cloning and Sequencing

A single primer set that would amplify relevant family members of the cannabinoid synthase clade was designed using the mapped cannabinoid synthase family from Genbank accession (SAMN06546749). Degenerate nucleotides were used at SNP positions within the 5’ and 3’ cannabinoid synthase coding sequence ends to capture the full-length genes present in genomic DNA of a given amplified variety (see Table 1A). The primer sets amplify gene targets of THCAS, CBDAS, CBCAS, and Clade C.

**Table 1.** A. THCAS Degenerate FL Primers and Slic Primers used for TOPO cloning, sequencing, and recombinant bacmid construct. B. List of *Cannabis* accessions screened used for TOPO cloning and sequencing. 120 clones were selected from PBBK accession, and 20 clones were screened from all other accessions. C. Randomly selected recombinant clones from Type III and Type II accessions, AC/DC and F-Cancer, for recombinant enzyme characterization.

| Accession | Clones |
| --- | --- |
| 1597/PBBK | 1-20;481-580 |
| ACDC 1147 | 381-400 |
| ACDC BK15575 | 401-420 |
| ACDC BK17861 | 21-40 |
| BK18700 Unknown Landrace | 61-80 |
| BK19139 Durban Poison | 41-60 |
| CAM4 | 81-100 |
| CAMH26 | 421-440 |
| Chinese 6 | 101-120 |
| Chinese 7 | 121-140 |
| Durban BK15526 | 141-160 |
| Super Lemon Haze | 161-180 |
| Durban Poison B | 181-200 |
| F-Cancer FLR1670 | 201-220 |
| FLR 1037 Durban Poison | 441-460 |
| Cannatonic 140620J011 | 221-240 |
| H2F | 241-260 |
| Harlequin 1946 | 261-280 |
| Hemp 2 Female | 281-300 |
| QC544-Super Goji OG Haze | 301-320 |
| Skunk 1 | 321-340 |
| HHC-744 BK17964 | 341-360 |
| Trigerion | 361-380 |
| UH18 | 461-480 |

**Table 1** A. THCAS Degenerate FL Primers and Slic Primers used for TOPO cloning, sequencing, and recombinant bacmid construct.
| Accession | Identity | C Tail Mod | Major Product |
| --- | --- | --- | --- |
| AC/DC | CBCAS | 6x His Tag | CBCA |
| AC/DC | CladeC.4 | 6x His Tag | CBCA |
| F-Cancer | CladeC.5 | 6x His Tag | CBCA* |
| F-Cancer | CladeC.6 | 6x His Tag | CBCA |

### TOPO Cloning and Sequencing

2.5uL of normalized gDNA was used as input into 22.5uL of Superfi PCR master mix prepared per reaction as follows: 12.5uL 2X Platinum^™^ SuperFi^™^ PCR Master Mix (Invitrogen), 1.25uL of 10uM forward and 1.25uL of 10uM reverse degenerate oligonucleotides (forward: 5’-ATGAAKTRCTCARCATTYTSYTTTTGGTWTG-3’; reverse: 5’-ATGAYGAYGYRGKGGAAGAGGTGGGAT-3’), and 7.5uL Nuclease free Water (Invitrogen). The reactions were subjected to the following thermocycler protocol: 1 cycle of 98C for 30secs; 35 cycles of 98C for 10 sec, 58C for 10secs, 72C for 45secs; 1 cycle of 72C for 5mins; 4C hold. End point PCR reactions were analyzed for target amplicon size bands at 1.6kb on a 1.5% agarose gel. The PCR product was used as input to carry out a TOPO cloning reaction and transformation with the TOPO^®^ TA Cloning^®^ Kit (Life Technologies) as per manufacturer instructions with TOP10 E. Coli competent cells. Analysis of transformants was carried out with X-gal blue/white screening on antibiotic selective media. For each transformation, 20-100 positive white colonies were picked for overnight culture in LB broth selective media and grown at 37C with shaking. Miniprep plasmid DNA isolation was carried out the positive cultures using an alkaline lysis procedure from the Plant Biotechnology Resource & Outreach Center molecular biology laboratory protocol (Reid, 1991). 580 Miniprep DNA samples were prepared for one premixed forward and one premixed reverse sequencing reaction with 1.6uM of M13 forward and M13 reverse sequencing primers and diluted in nuclease free water on a 96 well plate format. The premixed sequencing reactions were sent out for third-party sanger sequencing to Elim Biopharmaceuticals. Forward and reverse sanger sequencing reads were mapped to whole family of full-length cannabinoid synthases and percent identity match of read to each respective cannabinoid synthase was measured.

### Sf9 insect cell culture

Sf9 insect cells were purchased from Invitrogen (Waltham, MA), and initial p0 culture was thawed and recovered as per manufacturer instructions. The cells were maintained in T25 flasks with Sf-900^™^ II SFM and were grown in a 28C biological incubator removed from light exposure with aluminum foil coverings. Their morphology and growth patterns were observed with an inverted microscope and passaged 1:10 every 4 days once 95-98% uniform monolayer confluency was reached. Optimal transfections are carried out between 5-20 passages.

### SLIC subcloning and generation of recombinant bacmids for Cannabinoid Synthase Paralogs

2.5uL of normalized miniprep DNA was used as input into 22.5uL of Superfi PCR master mix prepared per reaction as follows: 12.5uL 2X Platinum^™^ SuperFi^™^ PCR Master Mix (Invitrogen), 1.25uL of 10uM forward and 1.25uL of 10uM reverse degenerate oligonucleotides with SLIC flanking homology to pFast donor vector and 6X His tag C terminal modification (forward: 5’-CGCGGATCCCGGTCCGAAGCGCGCGGAATTATGAAKTRCTCARCATTYTSYTTTTGGTWTG-3’; reverse: 5’-TACCGCATGCCTCGAGACTGCAGGCTCTAGTTAATGGTGATGGTGATGGTGATGATGAYGAYGYRGKGG AAGAGGTGGGAT-3’), and 7.5uL Nuclease free Water (Invitrogen). The reactions were subjected to the following thermocycler protocol: 1 cycle of 98C for 30secs; 35 cycles of 98C for 10 sec, 58C for 10secs, 72C for 45secs; 1 cycle of 72C for 5mins; 4C hold. PCR reactions were analyzed by diluting 1:2 in nuclease free water and 20ul was loaded into each well of a E-Gel^™^ EX Agarose Gels, 2%, 20gels and ran for 10 minutes on 1-2% gel settings for E-gel system. 1.6kb size amplicon was excised and purified using the SV gel clean up kit (Promega) and sent out for sequencing to confirm target. The pFast1 donor vector (Invitrogen) was linearized with BamH1-HF (NEB) and Pst1-HF (NEB) with CutSmart^®^buffer (NEB) under manufacturer conditions. A 10uL Sequence and ligation independent cloning (SLIC) reaction is prepared on the bench by mixing nuclease free water, a 1:1 molar ratio of linearized vector with purified SLIC PCR product (15ng-100ng of DNA total), 1uL T4 Polymerase (NEB), 1uL of 10X 2.1 NEB Buffer. The SLIC reaction is carried out for 5 minutes put onto ice to slow/stop enzymatic activity (Li and Elledge, 2012). 2.5uL of the Slic Reaction is used for subcloning to generate a recombinant donor pFast vector with the Bac-to-Bac^™^ Baculovirus Expression System (Gibco) as per manufacturer instructions using TOP10 E. Coli competent cells. Analysis of transformants was carried out on antibiotic selective media. For each transformation, 10-20 colonies were picked for overnight culture in LB broth selective media and grown at 37C with shaking. Miniprep plasmid DNA isolation was carried out the positive cultures using an alkaline lysis procedure from the Plant Biotechnology Resource & Outreach Center molecular biology laboratory protocol (Reid, 1991). The mini prep DNA was screened by PCR and positive clones were sent out for sanger sequencing as previously described for confirmation. The mini prep DNA was then used with the Bac-to-Bac^™^ expression system to generate a recombinant bacmid as per manufacturer conditions with MAX efficiency^™^ DH10Bac^™^ Competent Cells (ThermoFisher). After 4 hours of transformation recovery, cells were plated on LB agar with 50 μg/mL kanamycin, 7 μg/mL gentamicin, 10 μg/mL tetracycline, 100 μg/mL Bluo-gal, 40 μg/mL IPTG and incubated at 37C for 24-48h. Analysis of transformants was carried out with Bluo-gal blue/white screening on triple antibiotic selective media. For each transformation, 5-10 positive white colonies were picked and restreaked confirmation on selection plates to confirm white phenotype of colonies. Restreaked white colonies were picked for overnight culture in LB broth triple antibiotic selective media and grown at 37C with shaking. Recombinant bacmid DNA was isolated using the SV genomic DNA isolation kit (Promega) using manufacturer’s instructions.

### Sf9 transfection of Recombinant bacmid DNA and Recombinant Cannabinoid Synthase Expression

Purified bacmid DNA was prepared for quantification using the Qubit HS-dsDNA System (ThermoFisher) quantified using a Qubit Fluorometer (ThermoFisher) as per manufacturer’s instructions. Quantitated DNA was diluted to ~1ug/uL final working concentration and used as normalized input for bacmid transfections into sf9 insect cells. From 98% confluent and viable T25 flasks, sf9 cells were seeded into 6 well dishes at final ~8 x 10^5 cells per mL at a final volume of 3mL. A transfection reaction was prepared with ExpiFectamine^™^ Sf Transfection Reagent in 250 μL Opti-MEM^™^ I Reduced Serum Medium as described in the Bac-to-Bac^™^ expression system manual per manufacturer’s protocol (Gibco) for each clone in duplicate reactions. After incubation, the transfection mixture was added to each well. The cells incubated with the transfection mixture until a lytic morphological phenotype was observed indicating the generation of initial recombinant baculovirus product (P0), typically this occurred between 5-7 days post transfection. The supernatant from each well was collected and centrifuged at 300g for 5 mins once 90-100% lysis of transfected sf9 cells was observed. This supernatant contained the p0 recombinant cannabinoid synthase baculovirus which was removed from light and stored at 4C. 8 x 100uL of p0 supernatant was then added into 8 x T25 flask of 95-98% viable and confluent sf9 cells for each recombinant clone which generated ~80mL supernatant total in the final collection. Confluent sf9 cells incubated covered from light at 28C for 5-7 days and once a lytic phenotype was observed, the supernatant was collected. This subsequent infection generated a p1 stock of recombinant baculovirus and in the process target recombinant cannabinoid synthase was expressed and secreted from the sf9 cells into the supernatant. The supernatant was thereby collected from each flask and pooled to make 80mL supernatant total for each clone. This supernatant contained the 6x His C terminal tagged cannabinoid synthase protein, which was purified with nickel affinity column capture and purification.

### Ni+ Affinity Column Purification of Recombinant Cannabinoid Synthase

To purify each recombinant cannabinoid synthase, a single 5mL HisTrap^™^ Fast Flow column was used as per manufacturer guidelines with a flow rate of 5mL/min and 1 volume per 5mL (GE) for purification of each recombinant synthase. The following binding buffer was formulated: 100mM Tris, pH7, 15mM Imidazole and 250mM NaCl. Four Elution Buffers for step wise elution were formulated as follows: Elution 1) 100mM Tris, 250mM NaCl, 38mM Imidazole, Elution 2) 100mM Tris, 250mM NaCl, 65mM Imidazole, Elution 3, 100mM Tris, 250mM, 250mM Imidazole, and Elution 4) 100mM Tris, 250mM NaCl, and 350mM Imidazole and 5% Glycerol. A 60mL syringe attached to luer lock-capillary was used to create the flow and a manual 5mL/min plunging action with a digital lab timer was carried out. The column was equilibrated prior to use as per manufacture guidelines (GE). ~80mL of supernatant was supplemented with 15mM Imidazole and 100mM Tris final concentrations for Ni affinity column loading after column equilibration. Following equilibration, the supernatant was loaded onto the column. The column was then washed as per the manufacture guidelines. Following a column wash, the recombinant protein was eluted in with elution buffers 1-4 in a step wise fashion with 5mL volumes. Fractions collected using buffers 3 and 4 for each recombinant synthase were desalted using a PD-10 desalting column as per manufacturer instructions (GE). The purified protein was analyzed by SDS Page electrophoresis and western blot analysis using anti-His antibodies as per manufacturer guidelines (ThermoFisher).

### Cannabinoid Synthase Assay and HPLC analysis

Desalted protein extracts were quantified using a Qubit^™^ Protein Assay Kit and Qubit Fluorometer (ThermoFisher) as per manufacturer’s instructions. Enzymatic activity was measured in vitro in a reaction prepared as follows: 150uL of desalted purified protein extract 574uL 100mM Tris Buffer pH7 or 574uL 100mM sodium citrate Buffer pH4.5, and 26uL CBGA analytical standard (Restek). Reactions were carried out in sealed HPLC vials at 37C for 1 hour time increments. Reactions were stopped at various time points using 700uL MeOH. Reactions were filtered and loaded for HPLC analysis of proportion of CBGA converted enzymatically to other cannabinoids using an HPLC calibrated with known cannabinoid analytical standards. The data was acquired as previously described (Lynch et al., 2016).

### Promoter analysis

To examine the structure of cannabinoid synthase promoter regions we used tools from the meme suite to predict transcription factor binding sites. For a reference library of transcription factor binding motifs, we used the ArabidopsisDAPv1.meme file available at https://meme-suite.org/meme/doc/download.html. This is a DAP-seq data set representing about a third of known transcription factors in *Arabidopsis* (Bartlett et al., 2017). A cannabinoid promoter set was assembled by taking the thousand bases upstream of the start methionine for each gene, and these were searched with the DAP-seq motif database using fimo run with default settings (Grant et al., 2011). To reduce the number of false positive hits, a control set of 1000 random *Cannabis* promoters from the CS10 genome was prepared taking advantage of the reference annotation of gene structures to locate translation start sites. Next, we ran ame with --hit-lo-fraction 0.5 (McLeay and Bailey, 2010) to identify transcription factor binding sites that are specifically enriched in cannabinoid synthase promoters. Lastly, overlapping binding sites were resolved by selecting the highest scoring match.

### Assessment of amino acid conservation during gene family expansion

To investigate the nature of amino acid changes that occurred in cannabinoid synthase genes during the expansion of this gene family, we started by finding the best hit for THCAS, CBDAS and CBCAS from *Parasponia andersonii*, *Trema orientale*, and *Humulus lupulis*. For *Parasponia* and *Trema* this was done using the peptide data sets for these species. Such annotation does not yet exist for *Humulus*, so the most similar gene was found using blast, and then the gene model was refined using exonerate (Slater and Birney, 2005) so as to be sure to recover the full gene (which can be difficult with a tool like blast because sequence divergence, particularly in the signal peptide can be sufficient to stop the alignment before it reaches the start methionine). Interestingly, while these were the top hits with THCAS, the reciprocal did not hold true, with many of these hitting BBE genes down into the 66% identity range. This protein set was aligned using clustalw2, and the alignment is shown in (Supplementary Figure 11). Next the alignment was imported into R using the readAAMultipleAlignment function of the BioStrings package (Pagès et al., 2020). This was turned into a matrix, and then the severity of all amino acid changes was estimated using the Blosum62 substitution matrix as a scoring system (Henikoff and Henikoff, 1992). The control, or random, set was taken as the bottom diagonal of the substitution matrix, minus the perfect match scores. The two distributions were plotted using the R ggplot2 package (Wickham, 2016).

### Transcriptome Construction

The most complete collection of cannabinoid synthase genes, including both CBDAS and THCAS, is found in the Jamaican Lion Dash genome assembly (GenBank accession GCA_003660325.2), so many of the analyses in this paper were anchored to this assembly. As there is no annotation for this assembly, we created a reference guided transcriptome using Hisat2 version 2.1.0 and StringTie version 1.3.4d. RNASeq data used included trichome libraries from Zager et al. 2019. (BioProject PRJNA498707) comprising 27 libraries and the Cannbio2 tissue expression libraries from (Braich et al., 2019) (BioProject PRJNA560453) comprising 71 libraries. This data set is heavily weighted toward trichome data, but the Cannbio2 libraries bring substantial depth over a range of vegetative tissues. The bulk of *Cannabis* transcripts should be present in this set. Reads were aligned to Jamaican Lion using hisat2 with –max-intronlen 10000 and sorted bam files were created using samtools Version: 1.10 using htslib 1.10.2-3 (Li et al., 2009). Transcriptomes were computed tissue by tissue using StringTie with default parameters, and then the resulting set of output files was merged into a single transcriptome using the StringTie merge function. This resulted in 70,101 transcripts across 36,626 genes with a median length of 1552 bases. This is longer than the median length for the cs10 transcriptome (1140bp). Based on comparison of example sequences, it appears that the discrepancy comes from the StringTie transcripts being more likely to contain UTR sequence, or to contain longer UTRs. The difference in called gene number is probably just a byproduct of using the larger Jamaican Lion assembly.

### Gene expression

Publicly available rnaSEQ reads were mapped to reference genomes using Hisat2 version 2.0.6 (Pertea et al., 2016) setting a maximum intron size of 10kB and using the –dta option to generate output needed later in the pipeline for expression analysis. Next abundances were calculated also using StringTie using either our Jamaican Lion transcriptome or the NCBI cs10 annotation, depending on which reference genome was being used. Expression values as Fragments Per Kilobase Million (FPKM) were computed with ballgown version 2.26.0 (Frazee et al., 2015). Expression libraries used were the trichome libraries from(Zager et al., 2019) (PRJNA498707), the trichome libraries from (Booth et al., 2020) (PRJNA599437), and Cannbio2 tissue expression libraries from(Braich et al., 2019)(PRJNA560453). For Figure 2 the Braich and Zager trichome libraries were mapped to the CS10 assembly (GCF_900626175.1), while all other expression results shown are mapped to Jamaican Lion Dash (GCA_003660325.2).

## Results

### Expression of the *Cannabis BBE* family across tissues

To assess expression of the BBE family we used a cross tissue data set generated from Cannbio2 plants (Braich et al., 2019). Results are shown in Figure 1. This is a type II plant, so the THCAS and CBDAS genes dominate expression in trichomes and are much more lowly expressed in other tissues (although expression of these major cannabinoid synthase genes is never quite zero, even in root tissue). In female leaves CBCAS and THCAS account for most of the BBE family expression, while in male leaves a single clade of three unknown BBE genes accounts for most of the expression (Figure 1). More surprisingly, several of the uncharacterized BBE genes are also quite highly expressed in trichomes, although generally none of these appear to be trichome specific. While the cannabinoid genes have the least expression in root, many of the rest of the BBE genes show their highest expression in root. Lastly there are several of the BBE genes that show dramatically different expression in male versus female leaves, but without functional characterization there is no way to do more than speculate on what role this may be fulfilling.

**Figure 1.**
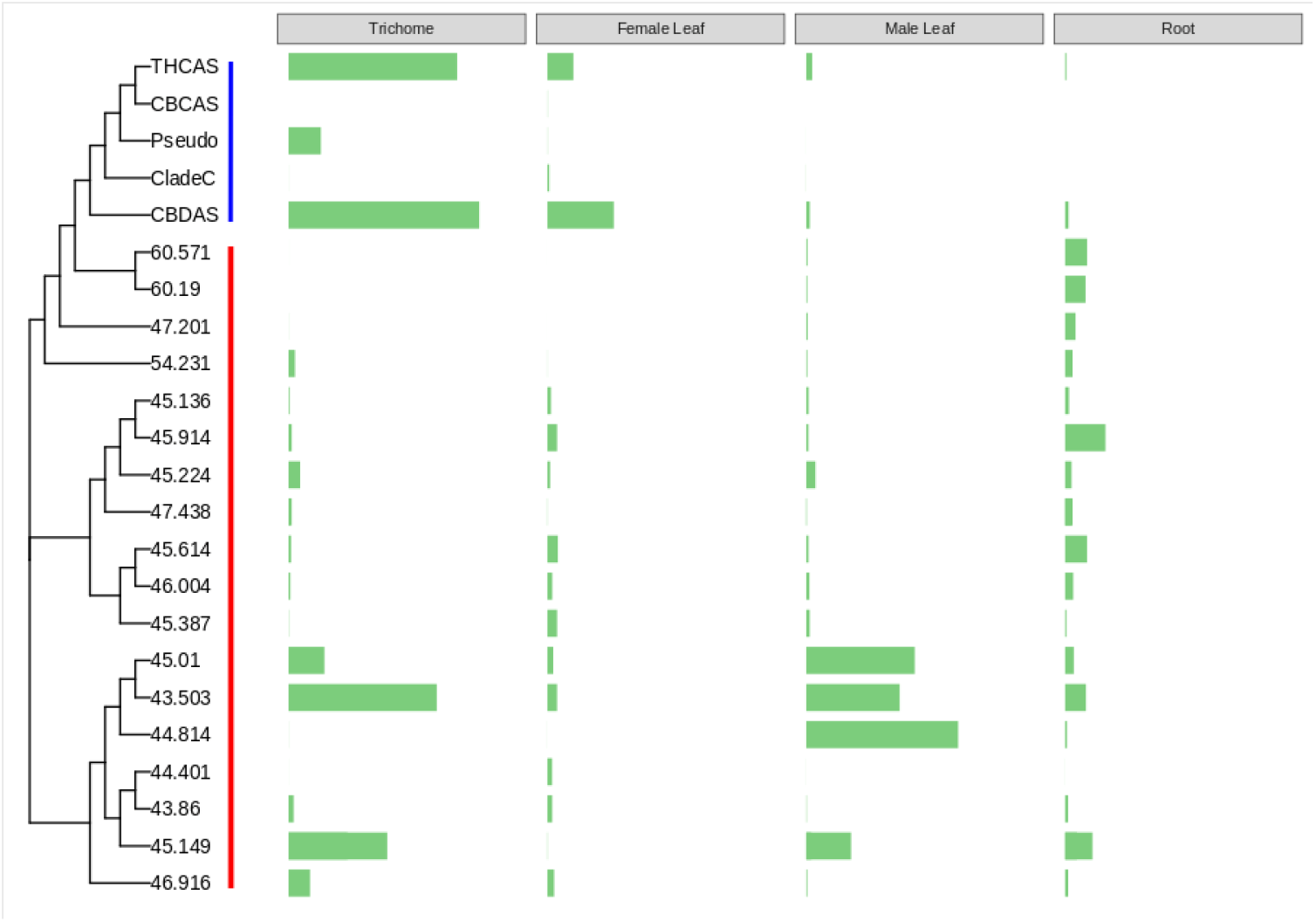
Expression of the BBE family across tissues. Tissue expression of the *Cannabis* berberine bridge enzyme family. On the left side of the panel is a neighbor joining tree of full-length protein sequences aligned using the clustalw2 algorithm (Larkin et al., 2007). The cannabinoid synthase group is highlighted with a blue line, while the rest of the uncharacterized BBE proteins are highlighted with a red line. The uncharacterized proteins are denoted with their percentage identity to the canonical THCAS protein sequence. The four bar plots show expression levels in trichomes, female leaf, male leaf and root tissue using the Cannbio2 rnaSEQ libraries (Braich et al., 2019). Bar height is percent total expression. Because the copies of the CBCAS and Clade C genes are so nearly identical, there is no reliable way to accurately determine individual expression values. Copies of CBCAS and Clade C genes are simply summed and collapsed in the figure. We have never observed more than one genomic copy of THCAS and CBDAS.

**Figure 2.**
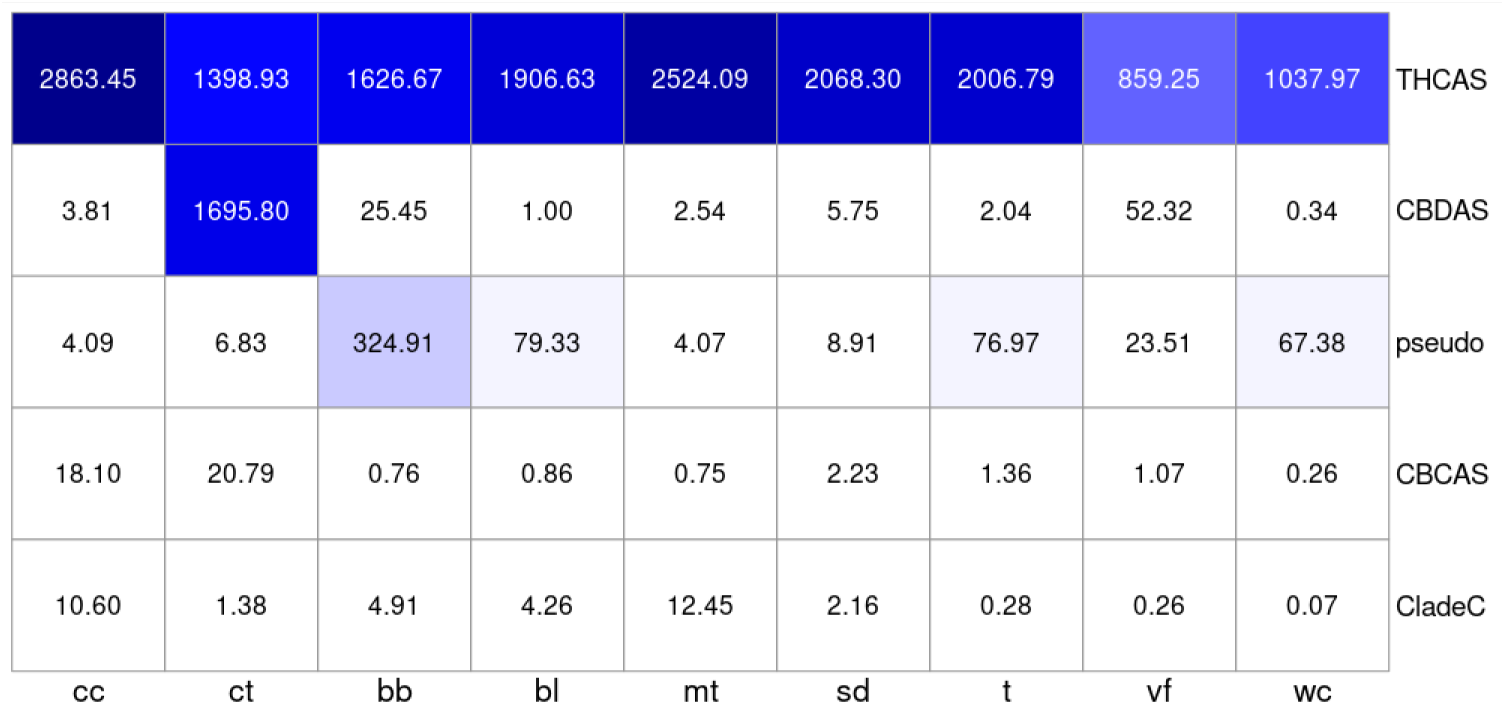
Trichome expression of cannabinoid synthase genes. Expression of cannabinoid synthase genes across trichome libraries. Strain abbreviations: Bb, Black Berry Kush; bl, Black Lime; ct, Canna Tsu; cc, Cherry Chem; mt, Mama Thai; sd, Sour Diesel; t, Terple; vf, Valley Fire; wc, White Cookies. Numbers shown are Fragments Per Kilobase of transcript per Million mapped reads (FPKM).

### Cannabinoid family gene expression in trichomes

To assess gene expression, we used a reference guided transcriptome built using the trichome expression libraries from (Zager et al., 2019), and the broad tissue expression libraries from (Braich et al., 2019). Jamaican Lion was used as reference genome because of its broader representation of genes in this family. For the heat map in Figure 2 just the nine trichome libraries from (Zager et al., 2019) are used. All but one of these strains is drug type, so not surprisingly THCAS expression dominates the pattern. CT is a type II strain and shows similar levels of THCAS and CBDAS. With both genes active, CT ends up with higher total cannabinoid synthase gene expression than any of the other strains. One of the issues to consider when evaluating expression in a family like this that has expanded by relatively recent gene duplication events, is that genes may be sufficiently similar to allow cross hybridization of 150 bp reads, complicating efforts to determine if any specific gene is expressed. We looked at the level of specificity for each gene by looking at which genes any multiply mapped reads map to. Most reads aligning to THCAS are uniquely mapped, but a small proportion also maps to the CBCAS genes and a smaller set that maps to the expressed pseudogene. CBDAS is sufficiently removed from the others at a sequence similarity level that essentially 100% of the reads mapping to it are mapping uniquely. The CBCAS genes are more difficult to approach. The six full length CBCAS genes from Jamaican Lion are separated by just 13 polymorphic sites across 1635bp. All reads mapping to a given gene in this set also maps to at least one other member of the CBCAS clade. On the other hand, most of the length of each of these genes is covered with reads that do not align to THCAS, CBDAS or the Clade C genes. While we cannot say exactly which of these genes are active, we can say that at least one of them is, and the observed expression is not spillover from other family members. Further, we can see that there is substantial variation in CBCAS expression levels across strains (although we don’t have data for notably high CBCA strains). The situation with the Clade C genes is similar, with the five full length Clade C genes in Jamaican Lion separated by 19 polymorphic sites across 1635bp. It’s clear at least one and probably several are expressed, but we don’t know which ones. Because of this, in Figure 2 the CBCAS and Clade C data are each combined into a single line.

Another interesting feature is what appears to be a highly differentially expressed pseudogene, classified as such because it lacks the first 35 amino acids, including the start methionine and the signal sequence, and contains two in frame stop codons starting at position 441 that remove critical parts of the enzyme active site. This means there is essentially no chance this transcript can make a functional product, but overall, it is the third most highly expressed member of this clade across these libraries, and in BB it is almost on a par with THCAS and CBDAS.

### TOPO Cloning and Sequencing

To survey genetic diversity of the highest homology targets in cannabinoid synthase family from a selection of a variety of *Cannabis* genotypes from the California recreational and medicinal market, a degenerate primer set was used to amplify target genomic DNA for full length cannabinoid synthase family members cloned into pET TOPO and subcloned with degenerate cannabinoid synthase primers modified with 6X His tag and pFastBac slic sites (Table 1A). The degenerate primer set was designed from the cannabinoid synthase family members with >85% protein identity to THCAS, targeting the canonical CBCAS but not CBDAS. Following PCR and TOPO cloning, 20 selected clones from each variety were sequenced. A previously sequenced genome PBBK (Steep Hill Labs, NCBI accession GCA_002090435.1) from which the degenerate primers were designed was used as a control and 120 clones were sequenced (Table 1B). The cannabinoid synthase family members most abundantly observed from the clone set were CladeC.5 and CladeC.6, which could be due to their prevalence in the cloned targets or could be a data artifact from the degenerate primer design as no targets for CBDAS were recovered (Supplemental Figure 11). Recombinant clones from Cannabis Type II and III chemotypes were assayed by recombinant enzymatic characterization (Table 1C). Chemotypes from selected libraries were measured by HPLC-PDA and cannabinoid quantitation (as previously described (Lynch et al., 2016; Vergara et al., 2017) (Figure 3A and 3B).

**Figure 3.**
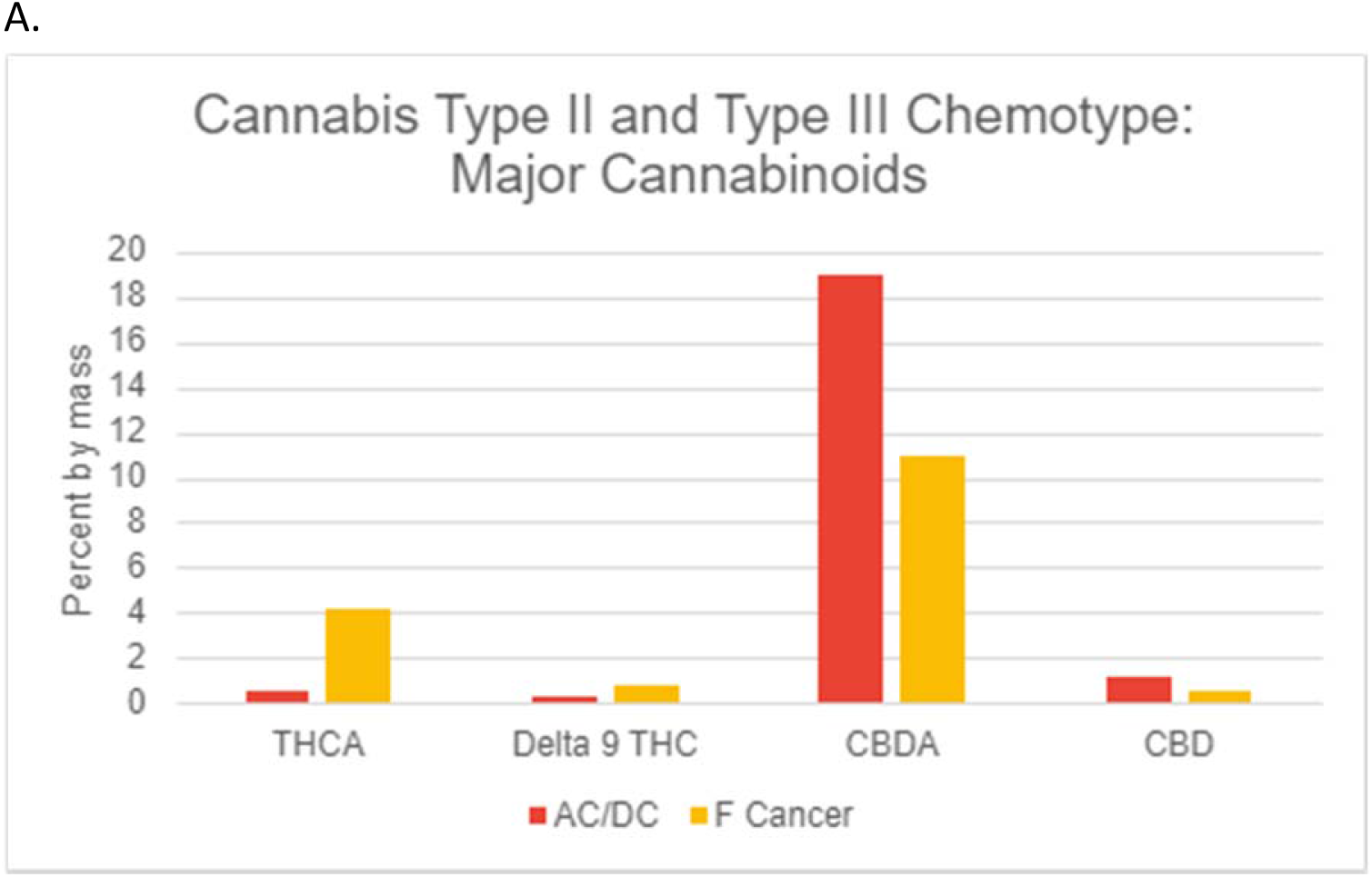
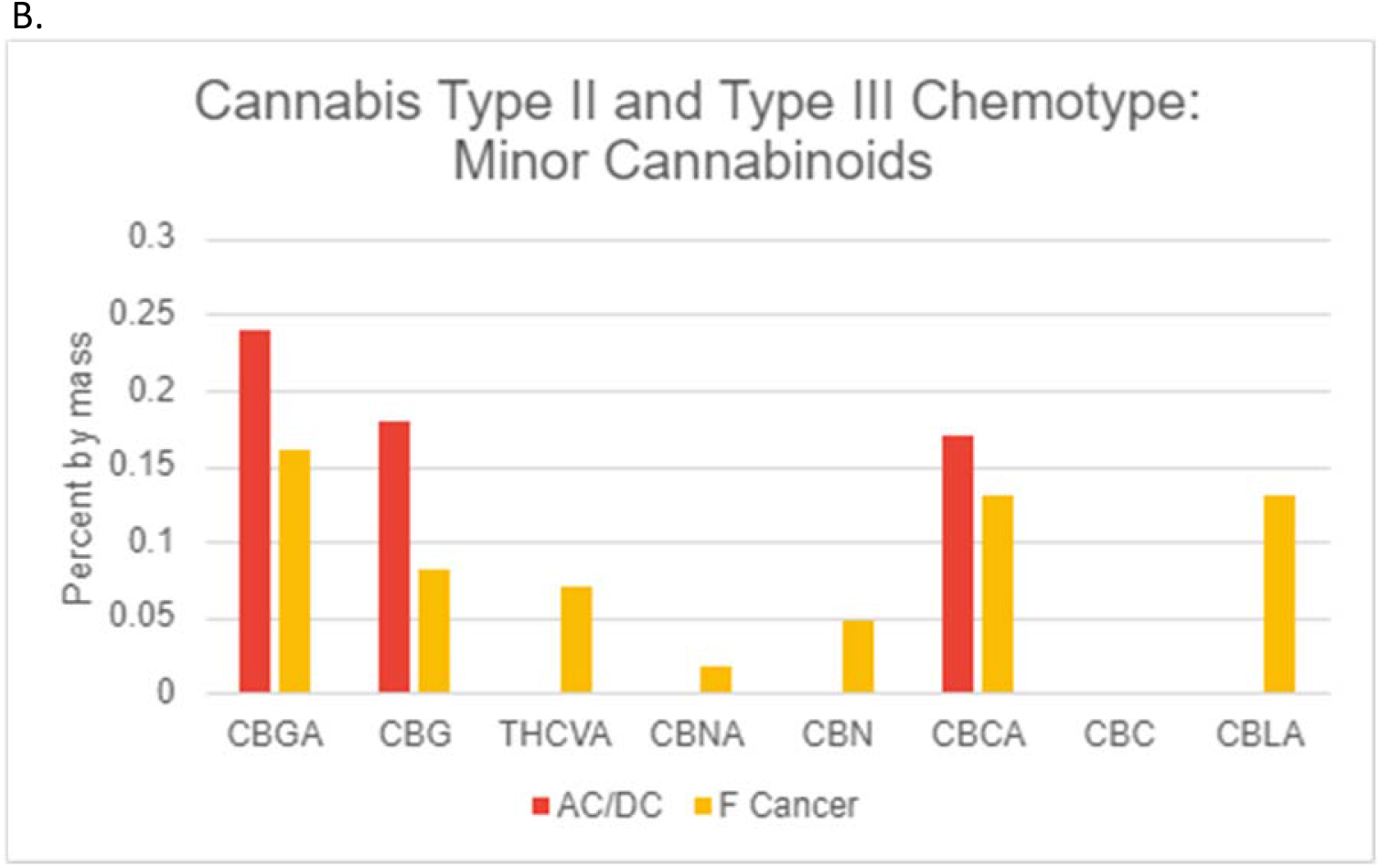
A and B Major and Minor Cannabinoid Chemotype from Type 2 and Type 3 CA Recreational *Cannabis* accessions.

### Cannabinoid Synthase In Vitro Enzymatic Characterization

To express the secreted form of each cannabinoid synthase, the plasmid constructs with native secreted signal peptide sequence and a 6X His tag terminal modification was subcloned into a Bac-to-Bac baculovirus expression (ThermoFisher, Waltham, MA) yielding a recombinant bacmid for transfection and subsequent infection in sf9 insect cells. Secreted cannabinoid synthase enzyme was collected/isolated/dialyzed and purified and analyzed by anti-his western blot analysis (Figure 4). His tagged enzyme products were sized at ^~^57KDa, which matches to previous reports (Morimoto et al., 1998; Laverty et al., 2019). 50-150uL of purified enzyme were used in vitro analysis with CBGA as a substrate and incubated under optimized conditions (Supplemental Figure 8). Notably, optimized chemical environment conditions for the invitro assays performed in this study were also similar to previous reports (Morimoto et al., 1998; Laverty et al., 2019). CBGA was used as precursor substrate and synthesized to CBCA product molecules. Post in vitro reactions were measured by HPLC-PDA absorbance detection at 258nm for CBCA (Supplemental Figure 3), 305nm for CBGA (Supplemental Figure 2), and 272nm for other side product cannabinoid acids observed in detectable amounts post incubation (Supplemental Figure 4-7). CBCA and CBCVA accumulation were also measured in CBGA and CBGVA substrate ratio dependent reactions (Supplemental Figure 9).

**Figure 4.**
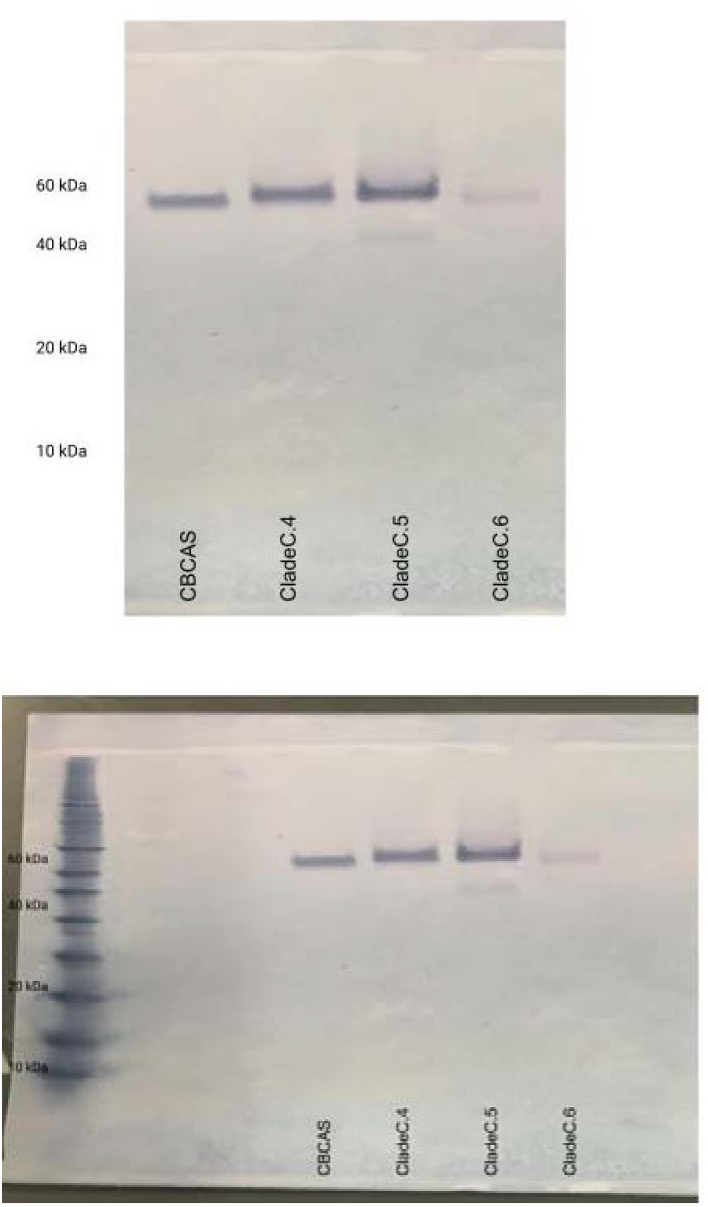
Anti-His Western Blot Analysis of Recombinant Cannabinoid Synthases. Anti-His immunodetection of recombinant protein at ~57Kda is observed in CBCAS, CladeC.4, CladeC.5, CladeC.6).

Five of the cannabinoid synthases characterized in this assay had CBCA synthase enzymatic activity as observed by the conversion of CBGA to CBCA as a dominant product (Figure 5). Percent CBCA conversion was carried out with area measurements of the precursor CBGA and product CBCA as previously described (Laverty et al., 2019). The reported canonical CBCAS denoted in this study as CBCAS and had a high homology matched with the reference CBCAS from (Laverty et al., 2019) (data not shown) was observed to have the highest level of activity among the cannabinoid synthases tested in this study. Moderate levels of CBCA were also observed in *in vitro* reactions from CladeC.5, and with lower activity in CladeC.4 and CladeC.6 a moderate amount of CBCA was observed following1h and 4h *in vitro* incubation (Figure 5).

**Figure 5.**
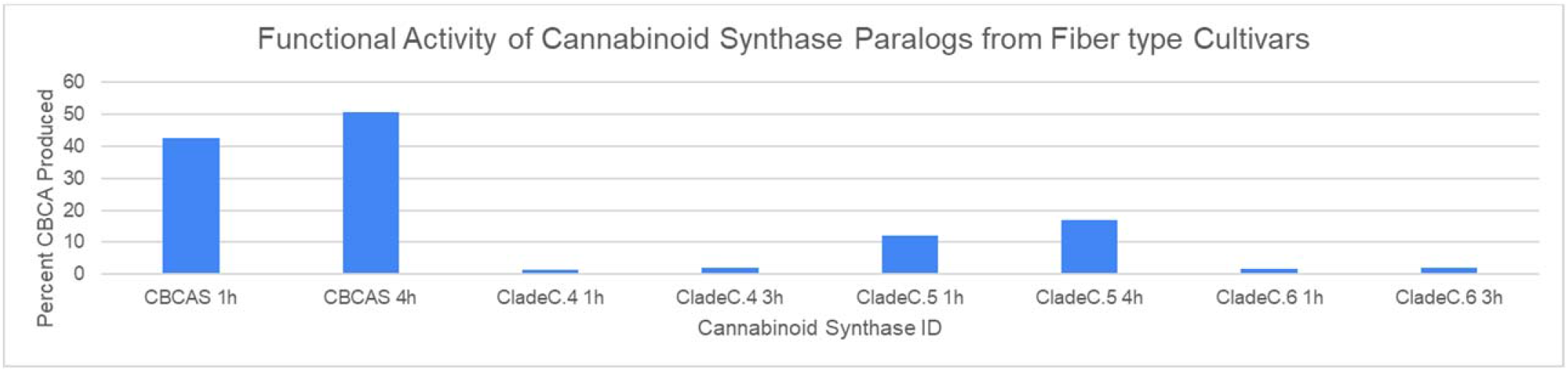
Functional Characterization of Recombinant cannabinoid synthases in CBGA supplemented invitro assay. Uniform protein and substrate concentration in CBGA Assay Buffer after 1h and 4h incubation at 40C. Activity presented as %CBCA produced with CBCA as the dominant product of these enzymes. Highest activity observed in the CBCAS identity recombinant enzyme. Lower activity of CBCA production was observed in CladeC recombinant enzymes.

To assess the kinetic differences between the putative CBCAS family, 50uL of purified enzyme was incubated with varying CBGA substrate concentrations and measured at specified time points with optimized reaction conditions following Michaelis-Menten principles previously described (Taura et al., 2007; Laverty et al., 2019). CBCA product velocity was plotted against substrate concentration in Graphpad Prism and IC50 Km and Vmax toolkit to determine Km, Vmax values from measurements and Kcat and Kcat/Km values were calculated all following Michalis-Menten input. Km values for CBCAS were similar to reported values (Morimoto et al., 1998; Laverty et al., 2019). Notably, Kcat and Km values of CladeC.5, CladeC.4, and CladeC.6 were observed to react with 2 orders of magnitude slower compared to canonical CBCAS (Laverty et al., 2019) CBCAS when performed under optimized assay conditions, however they were observably functional.

Figure 6 shows 15 residues shown in the crystal structure of (Shoyama et al., 2012) to be in the active site, or involved in binding the FAD cofactor molecule. The selected sequences span the diversity of the *Cannabis* cannabinoid synthase clade, in addition to the closest match from *Humulus lupulus*, *Parasponia andersonii* (PON38821.1) and *Trema orientale* (POO01102.1). As there is no annotation for the H. lupulus genome assembly (GCA_023660075.1) we identified the corresponding gene with a tblastn search against the genome assembly using the P. andesonii and T. orientale genes as queries. Of the 15 positions in this figure 7 are conserved all the way to *Trema* (note that the CladeC.5 and CladeC.6 sequences we obtained stop just short of the C-terminus, and so are missing these residues). Generally, these conserved residues have been shown to be necessary for function of the enzyme (Zirpel et al., 2018) As is the case with other members of the BBE family, the cannabinoid synthases have a covalently linked FAD moiety forming one side of the active site cleft (Shoyama et al., 2012). H114 and C176 are both covalently bound to FAD, and so it is not surprising that these residues are 100% conserved throughout this set. Y484 has been proposed to be the catalytic base that initiates the reaction by deprotonation of the hydroxyl group at the O6’ position of CBGA to make THCA (Shoyama et al., 2012), or deprotonation of a terminal methyl group of the geranyl residue of CBGA to produce CBDA (Taura et al., 2007). This residue is also completely conserved, including in the out species, suggesting that it is at least possible these non-*Cannabis* enzymes could share a similar reaction pathway. Another completely conserved position is L540, located on an N-terminal extension of the coding frame of either 8 or 16 amino acids. This extension has been found to be very important for enzyme function (Zirpel et al., 2018), to the extent that a stop codon at 539 almost completely abolishes activity. There are 11 positions that are conserved in the *Cannabis* enzymes but differ in the out species. One of these, Q124E substitutes an acid with an amide which, while considered a conservative change because both are polar, there could still be functional differences. Another of these, H292, which should be positively charged, is predicted to contribute to CBGA substrate binding by interacting with the negatively charged carboxyl group on CBGA (Shoyama et al., 2012). In the out species the histidine is replaced with arginine, a conservative change because the guanidino group on arginine will also carry a positive charge. Position A414 is also interesting because while the *Cannabis* enzymes have alanine at this position (except for CBDAS which substitutes valine), the out species have glutamine. Interestingly, Zirpel et al. (2018) showed that this change from alanine to glutamine was not only compatible with function but actually increased activity for CBDAS and THCAS. There are two positions that are conserved in the *Cannabis* enzymes and in *Humulus*, but differ in *Trema* and *Parasponia*. At position 175 the *Cannabis* and *Humulus* enzymes have a tyrosine, but this is replaced with valine in the *Parasponia* and *Trema* enzymes. This is a more substantial change, because this tyrosine residue is predicted to contribute to substrate binding by acting as a hydrogen bond donor (Shoyama et al., 2012). Tyrosine 417 also implicated in CBGA binding, is replaced with phenylalanine in *Trema* and *Parasponia*. While this is considered a conservative change, the hydroxyl group of Tyr 417 is predicted to participate in substrate binding as a hydrogen bond partner. H494 shows the most variability, and mostly tracks with clade. The CBCAS and CBDAS genes have a proline at this position, the THCAS genes have a histidine, and the clade C genes have phenylalanine.

**Figure 6.**
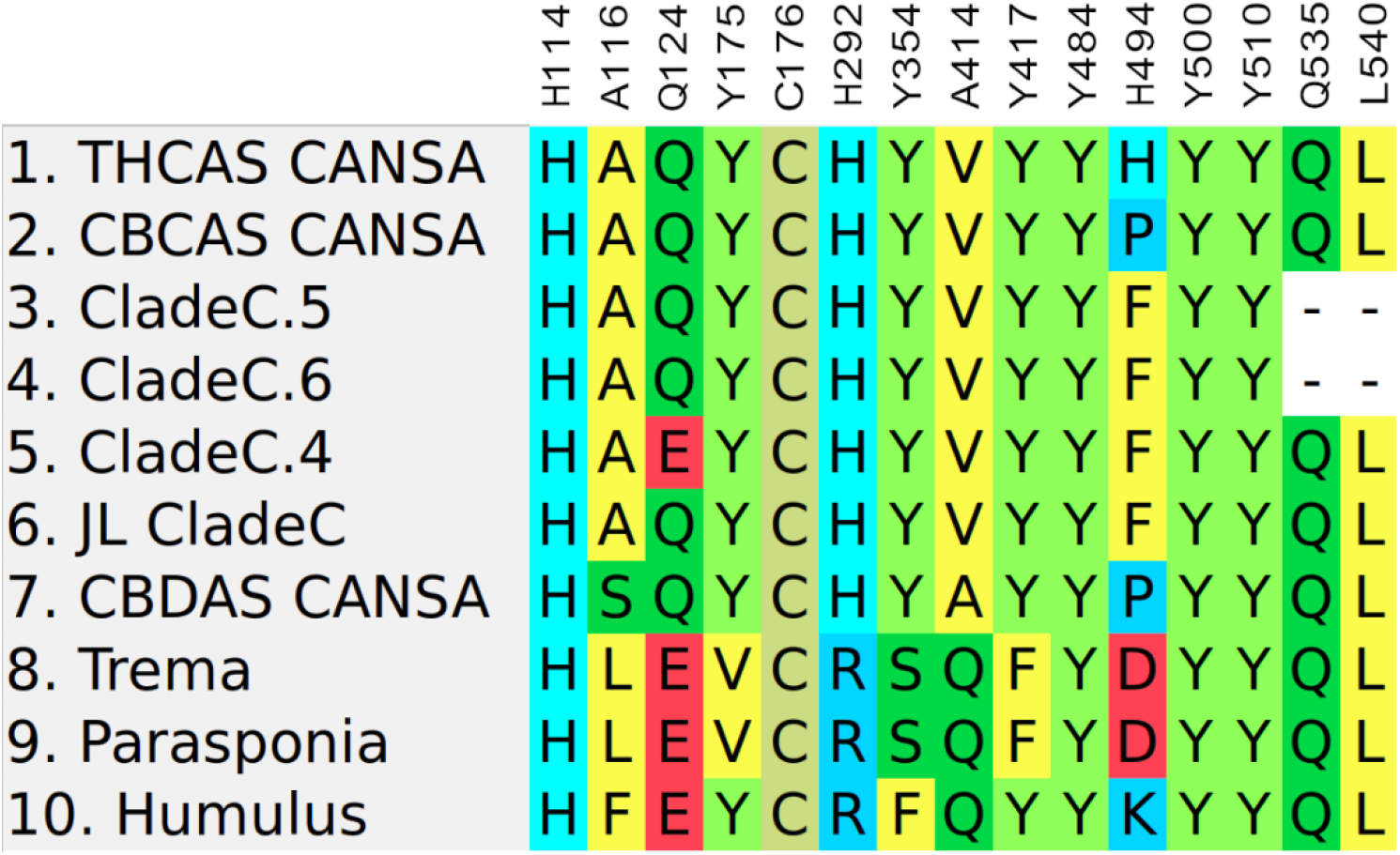
Active Site Residues. Alignment of residues that are in or near the cannabinoid synthase active site. Residue numbering consistent as previously described (Shoyama et al., 2012).

**Figure 7.**
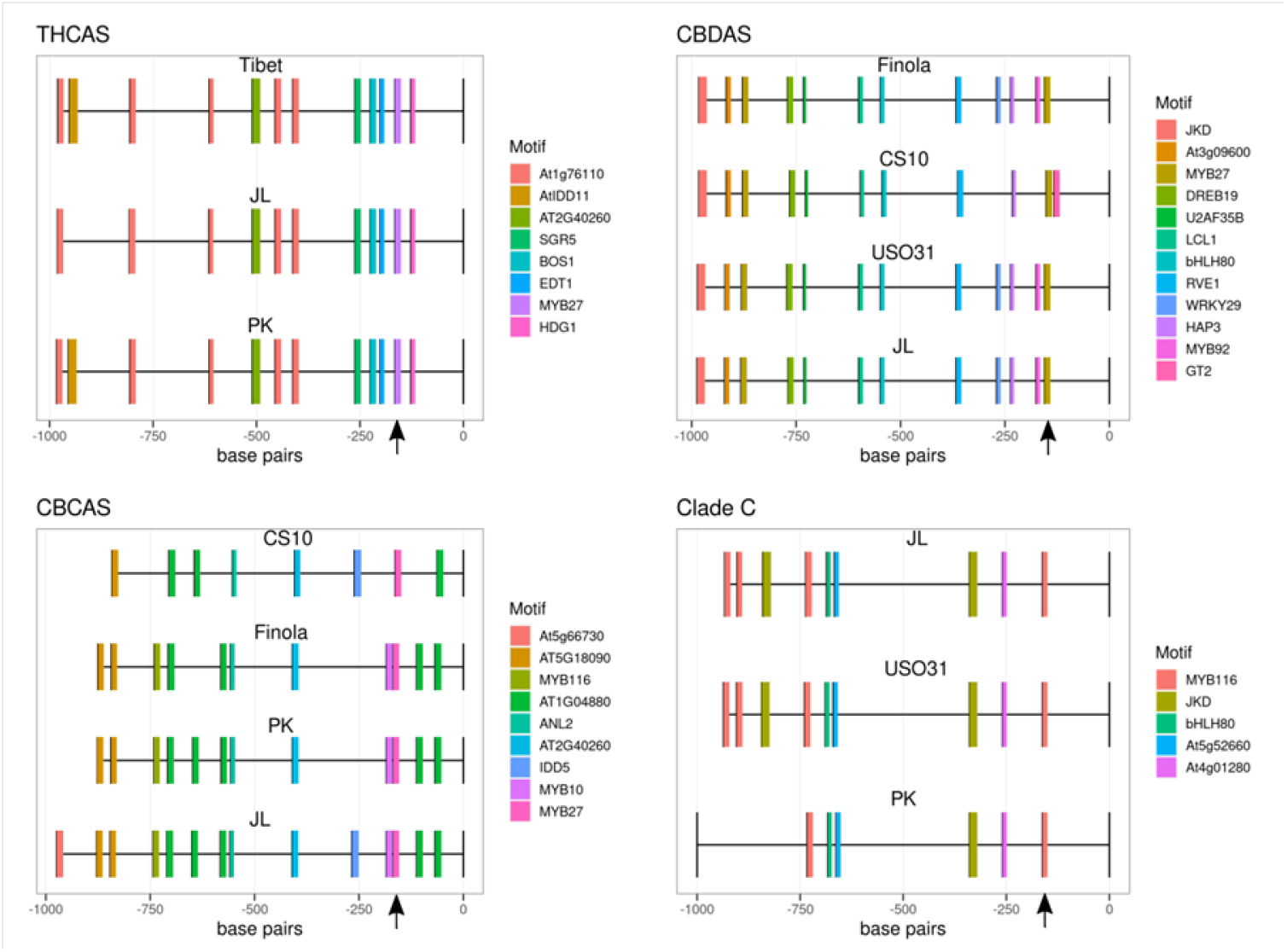
Predicted Promoter Structures. Predicted transcription factor binding sites from promoter regions for representatives of the four main *clades* of the cannabinoid synthase family. The genomes from which each was selected are shown. Cs10: reference genome (GCF_900626175.2), PK: Purple Kush (GCA_000230575.5). Finola: (GCA_003417725.2), JL: Jamaican Lion Dash (GCA_003660325.2), Tibet: JL (GCA_013030365.1), USO31 (manuscript in preparation).

### Analysis of promoter regions

Figure 7 shows the thousand bases upstream of the start methionine for representatives of the four cannabinoid clades from several different assemblies, with predicted transcription factor binding sites shown. The o-box identified as being necessary for promoter function is shown with a black arrow (Liu et al., 2021). The region from −227 to 0 is shared by all members of the cannabinoid synthase clade, diverging at about the same rate as the gene bodies themselves. That means that while the region is clearly shared, there is enough divergence that the predicted promoter binding sites between clades change quite a bit. In general, promoters for a specific family member are close to identical in all the assemblies we examined, but differences between clades are quite large. The o-box is identified in our analysis as a Myb27 binding site, rather than the AP2 binding site previously described (Liu et al., 2021). Figure 8 shows an alignment of the predicted Myb27 binding sites from representatives of the four clades, with the o-box indicated. There is adequate sequence divergence to at least suggest that binding properties to this site could differ between promoter types.

**Figure 8.**
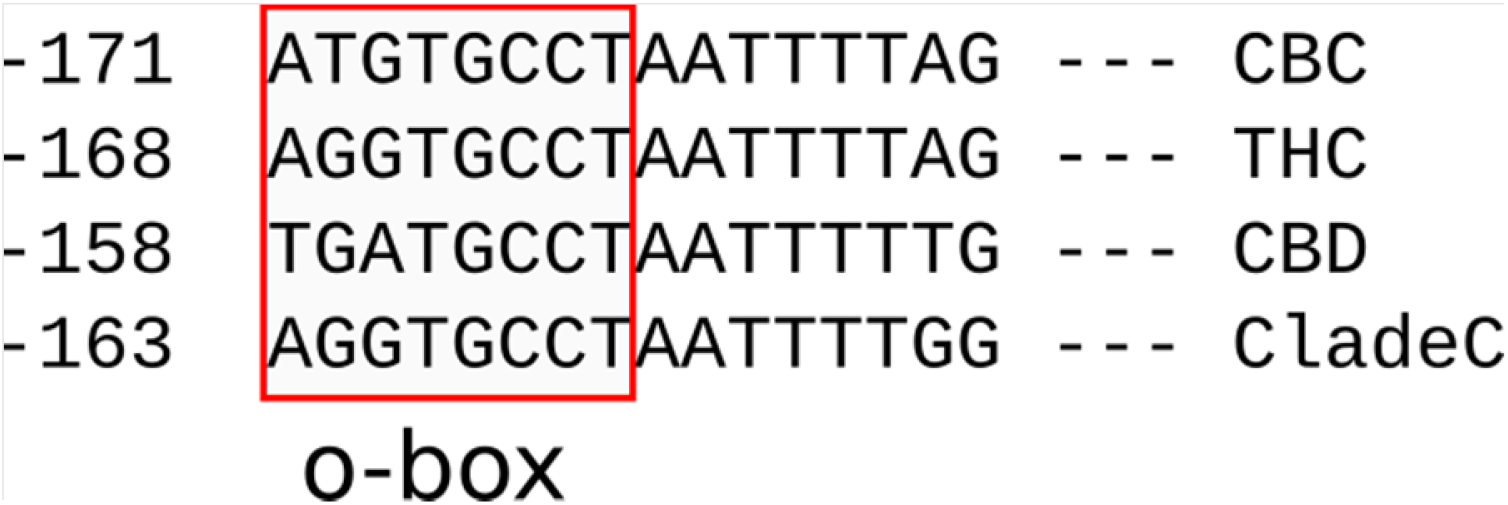
Alignment of o-box sequences. Predicted Myb27 binding site, containing the o-box (Liu et al., 2021) from representatives of the four cannabinoid clades from Jamaican Lion. The position of the leftmost residue, counting back from the start ATG, is shown along the left side.

A recent paper identified three transcription factors involved in cannabinoid synthase gene expression, csAP2L1, csMYB1, and csWRKY1 (Liu et al., 2021). In order to assess variation in the expression of these transcription factors in *Cannabis* trichomes we interrogated 14 trichome rnaSEQ data sets (Zager et al., 2019; Booth et al., 2020), and the results are shown in Figure 9. CsWRKY1 is represented by three variants called in the CS10 annotation, which change the coding sequence through use of an alternative splice site. CsMYB1 is far and away the most highly expressed of these three, but expression is variable across a 7-fold range. CsAP2L1 and csWRKY1 are more constant, varying across a 2 to less than 3-fold range, which is within the limits of experimental uncertainty. The ratio of csMYB1 to csAP2L1 is also quite variable, ranging from near parity (1.4 csMYB1:csAP2L1) to more than 7-fold.

**Figure 9.**
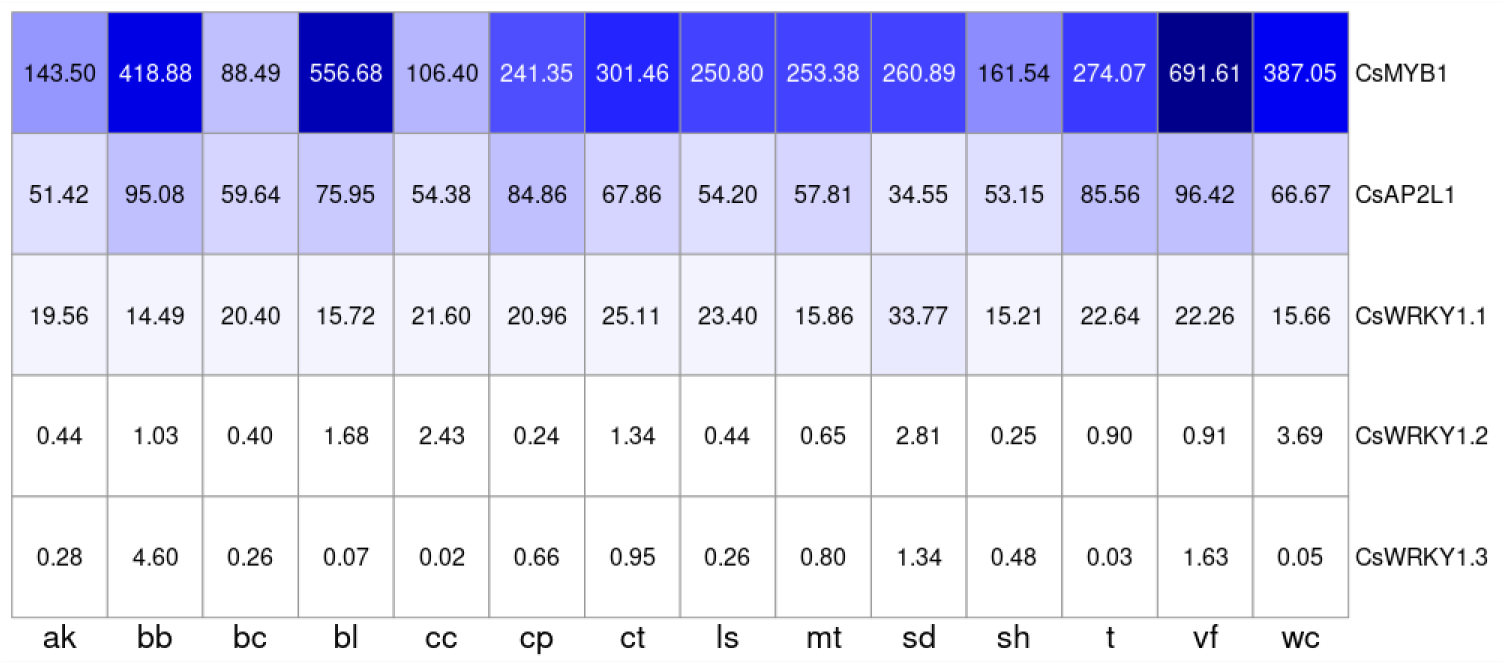
Expression of selected transcription factors in trichomes. Strain abbreviations: ak: Afghan Kush, bb: Black Berry Kush, bc: Blue Cheese, bl: Black Lime: ct: Canna Tsu, cc: Cherry Chem, cp: Chocolope, ls: Lemon Skunk, mt: Mama Thai, sd: Sour Diesel, sh: CBD Skunk Haze, t: Terple; vf: Valley Fire, wc: White Cookies. FPKM estimates are shown in individual cells.

### Shared promoter regions

All members of this family share an approximately 227 bp section upstream of the ATG start codon (denoted as region 1 in Figure 10), which includes the o-box region previously identified (Liu et al., 2021). As shown in the figure, the THCAS promoter shares a region extending out to about bp 298 with all members of the CBCAS cluster (region 2 in the figure), and this section of DNA is not seen in the other family members. This suggests that the genes in the CBCAS cluster originated from a duplication of the THCAS gene. Meanwhile, all members of the CBCAS cluster share a long region extending out to 1775 bp upstream of the start site, not seen in other clades (region 3 in the figure). This suggests that one product of the THCAS duplication went on to undergo a series of tandem duplications and given the very high degree of sequence similarity (>99%), this event was relatively recent. All members of the Clade C group share a longer segment extending out to 3618 bp upstream (region 4 in the figure). CBDAS, upstream of the region 1 sequence present in all these genes, has a segment extending out to about 640 bp that is shared with all the Clade C genes. Lastly, the highly expressed pseudogene only has part of this region, suggesting it arose from a different duplication event from the one that gave rise to the Clade C cluster. This work was complicated by the fact that we were running into sensitivity problems with blastn at about 85% identity, so the work was continued with discontiguous megablast, available as a task in the command line version of blastn. This provides a more sensitive search (at the expense of longer run times) and permitted us to see the rest of the shared regions.

**Figure 10.**
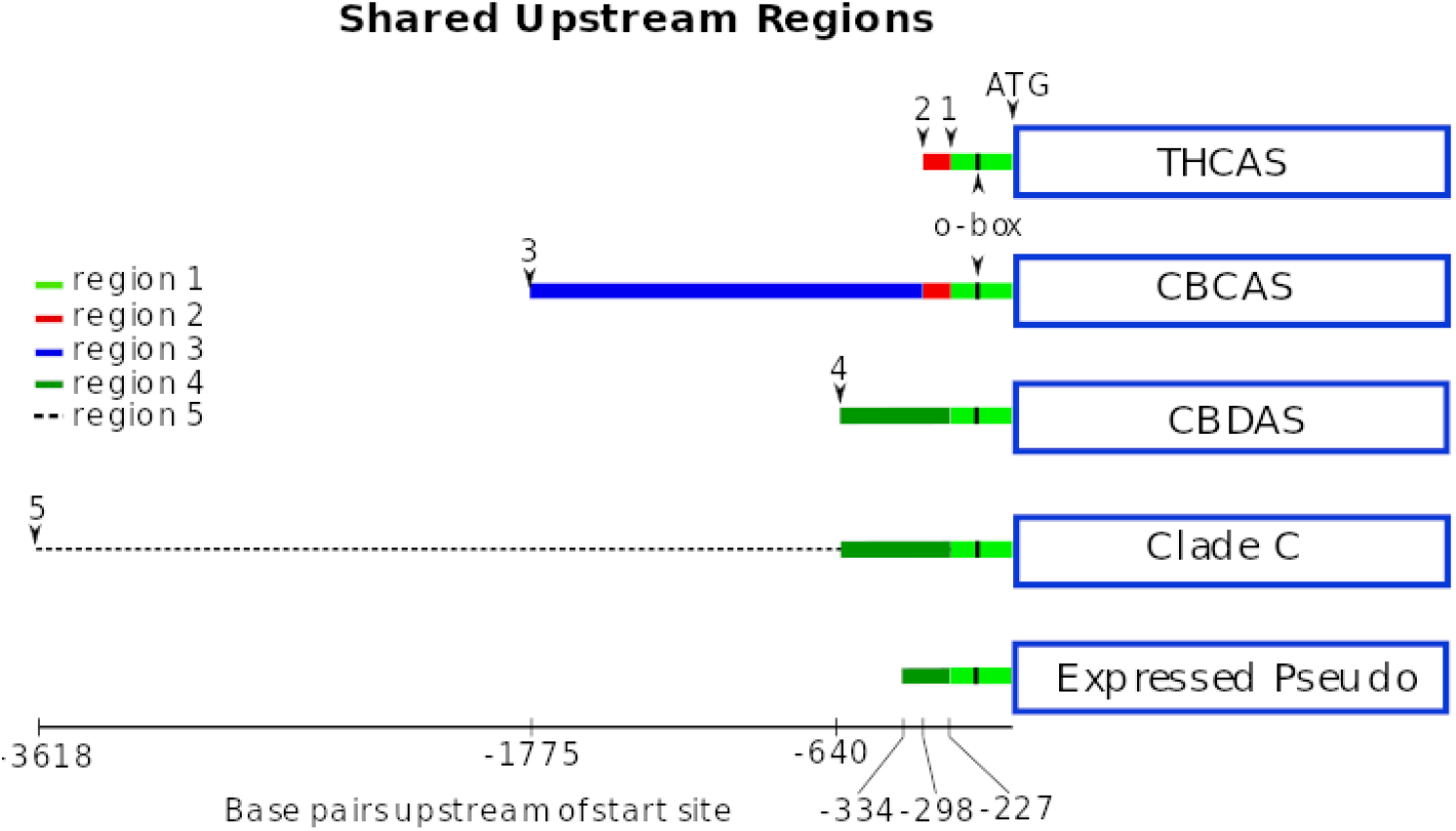
Shared upstream regions. Shared regions between upstream regions of cannabinoid synthase genes. Blue boxes indicate coding sequence, with the promoter region to the left, counting from the start ATG. Region 1, in green (extending to breakpoint 1), is shared by all members of this clade. Region 2, in red (extending to breakpoint 2) is shared by the CBCAS cluster and the THCAS gene (ie, all of clade A). Region 3, in blue (extending to breakpoint 3) is shared among all CBCAS genes, while Region 4 (black dotted line extending to breakpoint 4) is shared among all Clade C genes. Region 5 is shared by CBDAS and all the Clade C genes. The o-box identified by Liu, et al., 2021 is indicated with a black bar.

### Model to account for the pattern of shared promoter between cannabinoid genes

Figure 11 shows a model to account for the shared upstream regions in the cannabinoid family. Region 1, containing the previously identified o-box (Liu et al., 2021) is shared among all members of this clade, at the same break point, suggesting that this happened in a single event. Thus, we are positing that there is a progenitor gene that underwent tandem duplication giving rise to the THCAS, CBDAS genes and soon thereafter the CBDAS gene underwent another duplication to yield the progenitor to the Clade C genes. The initial progenitor does not seem to be still extant, and we were unable to find a relative within the broader BBE family that would be a candidate. THCAS, CBDAS and the Clade C genes are approximately the same number of mutations away from each other, as would be expected for three genes that arose around the same time from a common ancestor. Judging from the sequence similarities the next event in the expansion of this family was a series of tandem duplications by the Clade C precursor to yield as many as 8 genes (including pseudogenes), depending on the accession. More recently than that, the THCAS gene duplicated, creating the progenitor to the CBCAS cluster (which probably still exists as part of this cluster). The CBCAS cluster genes are very similar to each other, more so than the Clade C genes are to each other, indicating that the duplication event leading to the CBCAS cluster was more recent than that leading to Clade C.

**Figure 11.**
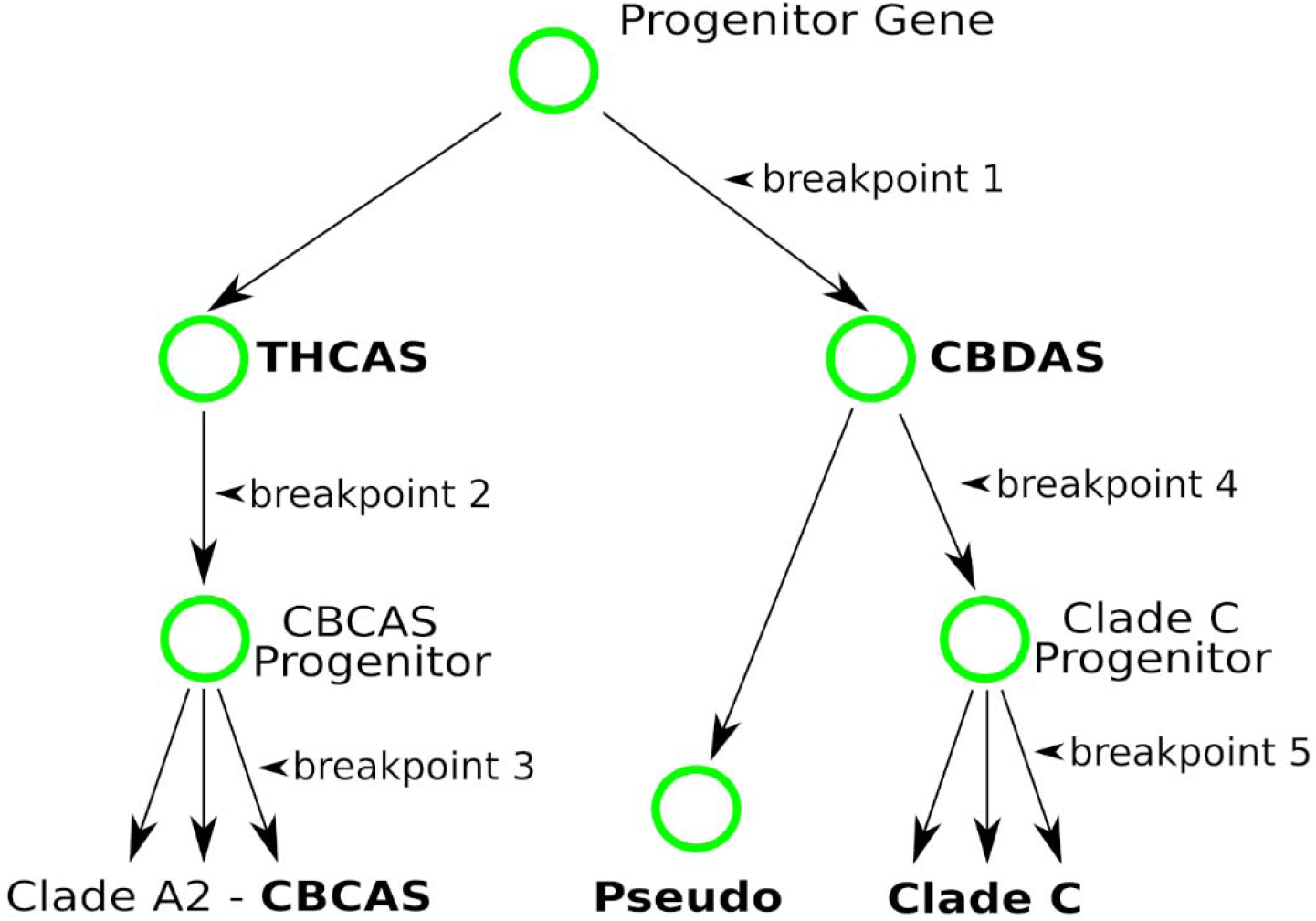
Proposed model for expansion of the cannabinoid synthase family. Proposed model of evolution of the cannabinoid synthase family, accounting for the pattern of shared regions in figure 10.

### Evolutionary distance of cannabinoid genes from related species

An easy way to visualize the differences between the BBE family in *Cannabis* and that in one of the related species is just to do a blast comparison and determine the distance (or number of mismatches) between each gene and its closest match in the out species. Figure 12 shows the result of this sort of comparison. First, each gene in the JL BBE set was compared to THCAS, and that similarity is shown on the x-axis (so THCAS itself is at bottom left, and CBDAS is to its left at 83% identity). The y-ais shows distance to the closest homolog in *Trema*. All the cannabinoid proteins are at a very nearly constant distance from the closest *Trema* protein, and they all have the same top match, PON38821.1, consistent with all these descending by duplication from a single gene as proposed by (van Velzen and Schranz, 2021). Interestingly, the rest of the *Cannabis* BBE family does not seem to have diverged nearly as much as the cannabinoid synthases, suggesting that there was selective pressure to maintain the function of these genes, while if anything there was pressure to innovate on the cannabinoid synthase clade.

**Figure 12.**
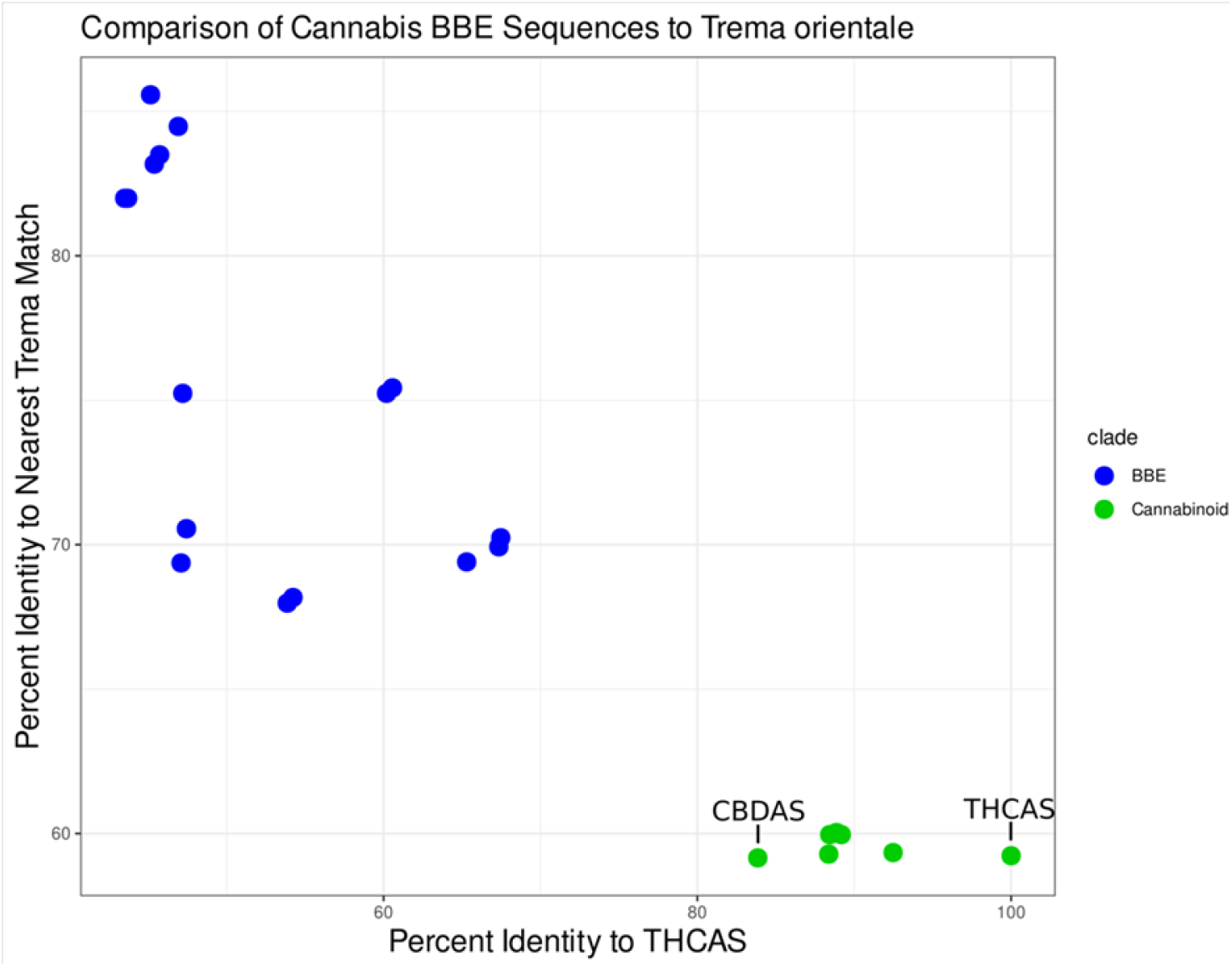
Comparison of the *Cannabis* BBE family compared to the *Trema* BBE family. Each point in the figure corresponds to an individual *Cannabis* gene, with the cannabinoid synthase clade in green and the rest of the BBE family in blue. The x axis is percent protein identity to THCAS, while the y axis is percent identity to the closest complete match in the *Trema orientale* protein data set.

### Evaluating amino acid changes during evolution of the cannabinoid synthase family

To get a view of the changes that occurred during expansion of the cannabinoid synthase family, we started by identifying the *Trema* BBE progenitor gene. Figure 4a in (van Velzen and Schranz, 2021) shows the syntenic region in *Trema* residing at one end of contig JXTC01000120.1, but the best protein hit to THCAS resides on a different contig, JXTC01000010.1. Closer examination shows that the *Trema* genome contains 35 full length BBE genes, with 28 in a large cluster at one end of JXTC01000010.1, another five at one end of JXTC01000120.1, and two more as singletons on other contigs. We further found that the BBE gene cluster on JXTC01000010.1 is bounded by several cationic amino acid transporter genes, which also bound that end of the corresponding cluster in Jamaican Lion. This suggests that these two contigs are adjacent on the chromosome, together constituting the syntenic site identified (van Velzen and Schranz, 2021). Given that the BBE genes on JXTC01000120.1 are much more distantly related to THCAS, we selected the best hit, which is several genes further into the cluster on JXTC01000010.1. Next the corresponding gene from *Parasponia* and *Humulus* was identified by homology. The protein sequence for these three genes were aligned using clustalw2 to representatives of the cannabinoid synthase family (Supplementary Figure 11). The alignment was imported into R, and every amino acid change in the alignment was assigned a conservation score using the BLOSUM62 scoring matrix. This was compared to random by taking all the non-match scores from the matrix. Results are shown in Figure 13 Conservative changes (like isoleucine to valine) get higher scores, and non-conservative changes get negative scores. The scores for random changes skew quite heavily into negative territory, with a mean score of −1.53, but more importantly, 78% of the random changes had scores less than zero, and just 11% had positive scores. On the other hand, the total set of changes that have occurred during evolution of the cannabinoid family tend much more heavily into positive terrain, with a mean very close to zero, and only 40% of changes having scores less than zero and 37% positive. This suggests that during evolution amino acid changes have strongly tended to favor changes that more closely maintained function of the source non-*Cannabis* gene.

**Figure 13.**
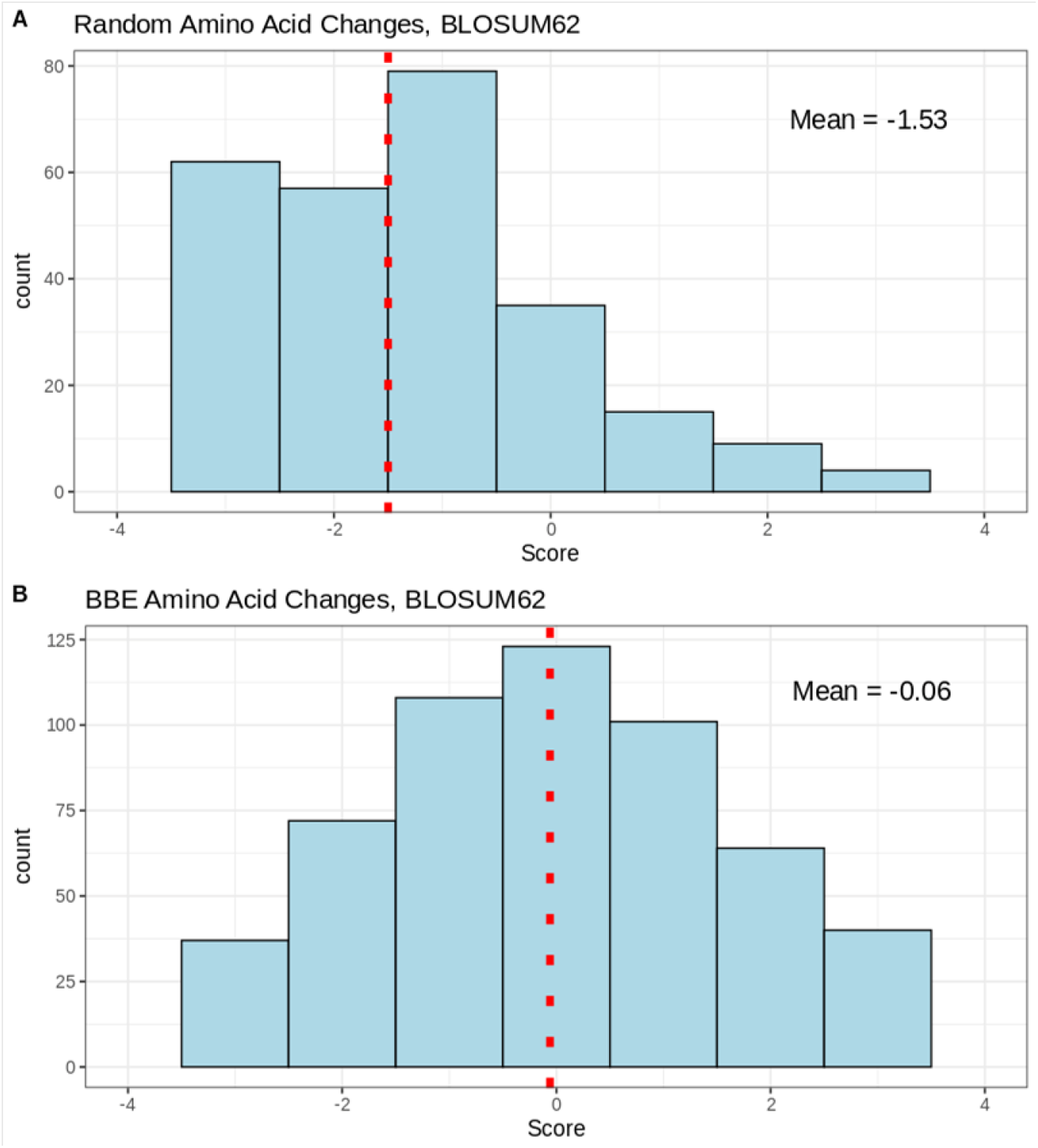
Predicted effect of amino acid changes during expansion of the cannabinoid synthase family. Nature of amino acid substitutions during evolution of the cannabinoid synthase family. A. Full set of substitution scores from the Blosum62 matrix. B. Blosum62 scores for substitutions that have occurred during evolution of the cannabinoid synthase family.

## Discussion

Several features of the cannabinoid synthase family have made it difficult to accurately describe in detail. In the early investigations aimed at understanding cannabinoid biosynthesis, it was unclear that this was a gene family with up to a dozen closely related genes. In particular, the CBCAS genes are very similar to THCAS, with 92% identity at the protein level, and 96% identity at the DNA level. This high similarity has led to some confusion in the literature (Hurgobin et al., 2021), and in fact many of the CBCAS genes currently in GenBank are annotated as THCAS. A further issue is the extent of strain-to-strain variation in *Cannabis*, complicating the task of judging whether a new sequence represent allelic variation, which is quite high in *Cannabis*, or a separate gene. Another issue is that presence absence variation (PAV) is substantial, but primarily among the A2 (CBCAS) and C clades. We agree with others who have concluded that there do not seem to be strains that contain more than a single THCAS gene, or a single CBDAS gene (Hurgobin et al., 2021; van Velzen and Schranz, 2021; Thomas and Kayser, 2022). The THCAS and CBDAS genes are tightly linked in a region that appears to have suppressed recombination (Kojoma et al., 2006), and also appear to be in repulsion (Toth, et al., 2019). An interesting note about this region is that of the two genome assemblies we have for type II *Cannabis*, Cannbio2 and Jamaican Lion, the first only includes THCAS in the assembly, while the second assembles both genes but places them on different large contigs. Both strains have fully functional copies of both genes. In comparing the Type I Purple Kush with Type III Finola, (Laverty et al., 2019) found that the chromosomal segment containing CBDAS in Finola had almost no similarity to the Purple Kush segment containing THCAS. This would support the idea that Type II plants have these genes in repulsion rather than on the same chromosome, but further work will be required to fully understand the B locus.

Cannabinoid synthase expression in trichomes is overwhelmingly dominated by THCAS and CBDAS, with expression levels typically several orders of magnitude higher than the CBCAS and Clade C genes. The CBCAS genes are difficult to assess because their similarity to THCAS is great enough that a small subset of rnaSEQ reads that align to THCAS will also align to a CBCAS gene, meaning one can’t say for sure where these reads originated. Fortunately, most of the reads aligning to CBCAS genes are not shared with THCAS, and vice versa, even if they are shared across the entire CBCAS clade. We can say with confidence that at least one of the CBCAS genes is active in most of the strains examined here, and that there is substantial variation across strains for CBCAS expression, but the similarity within the CBCAS cluster is so high that it would be very difficult to say if it’s only one or if all the intact CBCAS genes are active. The Clade C genes are less problematic given they have a slightly higher sequence divergence from the other clades, meaning reads mapping to a Clade C gene will not map to genes in the other clades, but again, similarity within this group is almost as high as within the CBCAS clade making it is difficult to know if one or all of them are active.

Looking at expression across tissues using the Cannbio2 expression set (Braich et al., 2019) the first important point is the amount of expression happening across the *Cannabis* BBE family, including several completely uncharacterized enzymes that are expressed in trichomes at levels comparable to the dominant cannabinoid synthases. The BBE genes outside the cannabinoid clade are expressed throughout the plant, including in root tissue. A number of these genes also show marked differences between male and female leaf, with several appearing to be nearly or even completely male-specific.

One of the goals of this study was to increase the number of characterized genes in this family. To this end, we selected a subset of the family by using degenerate primers that amplify targets > 85% protein level identity to THCAS, using genomic DNA isolated from broad sample set of accessions of type 1, 2, and 3 (de Meijer et al., 2003) commercial *Cannabis* varieties from California. PCR and cloning yielded about 500 recombinant clones of various cannabinoid synthases from the germplasm pool. Due to limited resources, we chose to prepare recombinant baculoviruses from 7 isolates with 88-92% protein level identity to THCAS and perform infection/recombinant expression in sf9 insect models. (Laverty et al., 2019)(Morimoto et al., 1998; Laverty et al., 2019) The other 3 novel active synthases had 88.8-89.2% protein level identity to THCAS and were identified as Clade C genes using the previously proposed nomenclature (van Velzen and Schranz, 2021). We also found that the Clade C genes characterized here also make CBCA as primary product, and while there are side products, there was no THCA detected. (Thomas and Kayser, 2022) showed that CBCAS activity with no THCA or other side products detected and that THCA and CBCA were side products of CBDAS. (Fulvio et al., 2021) examined CBCAS genes from several different accessions, and while these genes are almost completely identical, they did find 28 sites where mutations led to amino acid changes. Seven of these sites are in the first 28 amino acids, which is the predicted signal peptide, and hence are unlikely to affect function. The rest of the sites are distributed across the mature protein sequence, and while none of them are in the predicted active site, it is possible one or more of these could change product profile. Nonetheless failure to find THCA as a side product of the CBCAS enzyme makes it less likely this is the source of THCA in strains that lack a functional THCA synthase. Further, the poor kinetic properties of the Clade C genes, with kcat values as much as two orders of magnitude lower than the canonical CBCAS enzyme (Table 2). Combined with the low expression seen for these genes (Figure 2) it seems unlikely they contribute in a significant way to oil composition, although there may well be varieties yet to be tested with higher levels of expression for the Clade C genes. Finally, the ratio of CBD:THC typically found in lines lacking a functional THCAS gene (around 24-28 to 1) is close to what has been measured for the pure enzyme at its pH optimum of 4.5 where THCA and CBCA are each produced at about 5% of total cannabinoids (Zirpel et al., 2018), indicating that the CBDAS enzyme itself is the source of this “extra” THCA.

**Table 2.**
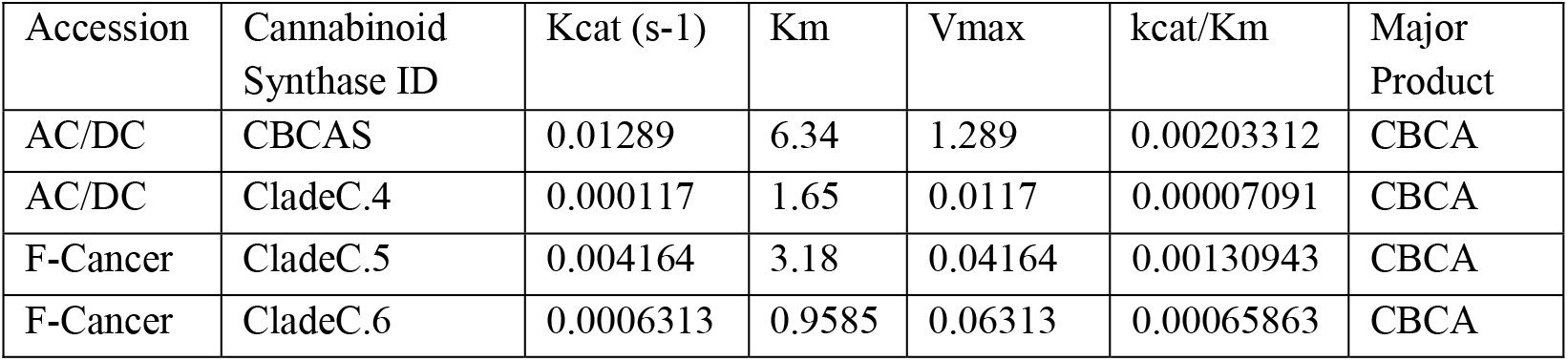
Kinetic Values for Recombinant Cannabinoid Synthases CBCA product velocity (y axis) was plotted against substrate concentration level (x axis) in Graphpad Prism and IC50 Km and Vmax toolkit to determine Km, Vmax values from measurements and Kcat and Kcat/Km values for recombinant enzymes in the CBGA *in vitro* assay. Experimental value kinetics were calculated all following Michalis-Menten input with normalized protein concentration in increasing substrate concentration.

Molecular clock techniques have been used to estimate that *Cannabis* and *Humulus* diverged from each other 27.8 Ma (Ren et al., 2021), and the earliest fossil pollen consistent with *Cannabis* is from 19.6 Ma, found in NE China (McPartland et al., 2019). Large scale resequencing efforts across broad germplasm shows evidence of human selection pressure on Cannabis for something on the order of 10,000 years, an estimate that agrees well with fossil evidence of *Cannabis* use by neolithic populations (McPartland et al., 2019). Our analysis of the amino acid changes that have occurred during expansion of this family also show evidence of selection.

The model we are proposing to account for the shared upstream regions in the cannabinoid synthase family is broadly similar to that proposed by (van Velzen and Schranz, 2021), with the exception that we see the separation of THCAS, CBDAS and Clade C as two steps with the Clade C progenitor arising from a duplication of CBDAS. But this second step would have occurred soon after the THCAS/CBDAS split since all these genes are about the same distance from each other. Further, the entire family is very close to the same similarity distance from the closest counterpart in *Trema*, which is what would be expected if, consistent with results from Weiblen, et al., 2015 this family evolved from a single progenitor gene that existed prior to *Cannabis* speciation.

The promoter analysis work reported here falls closely in line with the evolutionary model in that shared promoter regions are diverging at about the same rate as the gene bodies. Within a clade the promoter structure is quite similar, in particular between CBCAS genes, which appear to be a very recent series of duplications, and somewhat more variable in the Clade C genes, which look like an older duplication, but it’s very different between clades. The only region shared by all members of this family is the o-box identified by (Liu et al., 2021), which is necessary for CsAP2L1-induced transcriptional activation of THCAS promoter. Our computational prediction analysis identified this element as a MYB27 binding site, but this is based on similarity to described binding sites in *Arabidopsis*, and so cannot necessarily identify the precise transcription factor.

Given that the progenitor of the cannabinoid synthases is found in the related genera *Humulus, Parasponia* and *Trema*, it is reasonable to ask whether these enzymes actually make cannabinoids. In fact, it has recently been found that THCA, CBDA and cannabinol (CBN) are present in inflorescences of *Trema orientalis* (Napiroon et al., 2021). Looking at the active site residue alignment in Figure 6 the *Trema* sequence, which is about 60% identical to the *Cannabis* cannabinoid synthase sequences shows about the same percentage identity in the active site. Many of the differences are in positions that are also variable among the *Cannabis* genes, but there are four changes at sites that do not vary in the *Cannabis* genes, including a Tyr → Val change that eliminates a hydrogen bond donor implicated in substrate binding. Nonetheless, the *Trema* gene could well be a cannabinoid synthase, and there are probably other candidates among the *Trema* BBE family that could account for the observed cannabinoid production.

The expansion of the cannabinoid synthase family in *Cannabis*, seemingly all from a single progenitor gene after speciation, is a remarkable event in the history of the BBE gene family. That this expansion is ongoing is evident from the recency of the gene duplication events that gave rise to the CBCAS (Clade A2) and Clade C genes, and the fact that copy number of these genes is highly variable between strains. Clade C genes primarily make CBCA, along with a few side products that were not identifiable using currently available *Cannabis* chromatography standards (see Figs. S3-S7), indicating that evolution continues to explore the biosynthetic potential of this family. The Clade C genes in the Jamaican Lion assembly have just 19 polymorphic sites across 5 genes but given the high level of variability among *Cannabis* accessions, it is fair to say that the variation among Clade C genes has just barely been sampled. The fact that Clade C genes predominantly make CBCAS does narrow the biosynthetic options a little given that these enzymes are specialized to make an already relatively abundant product (CBCA) rather than minor cannabinoids. But these compounds will occur through alternative substrates (as with the varins) or by subsequent modification of the primary cannabinoids.

## Acknowledgements

The authors would like to thank Daniella Vergara and Robert Givens for useful discussions, and Christopher Pauli for assistance with the manuscript.

## Author Contributions

KDA, AT, and RG conceived and planned the work. AT performed all of the molecular biology, including the enzyme assays. KDA performed the bioinformatics analysis. KDA and AT wrote the manuscript, KdC contributed to the interpretation of the results, and all authors took part in the editing of the manuscript.

## Parsed Citations

**van Bakel H, Stout JM, Cote AG, Tallon CM, Sharpe AG, Hughes TR, Page JE (2011) The draft genome and transcriptome of Cannabis sativa. Genome Biol 12: R102** Google Scholar: <u>Author Only Title Only Author and Title</u>

**Bartlett A, O’Malley RC, Huang SC, Galli M, Nery JR, Gallavotti A, Ecker JR (2017) Mapping genome-wide transcription-factor binding sites using DAP-seq. Nat Protoc 12: 1659–1672** Google Scholar: <u>Author Only Title Only Author and Title</u>

**Booth JK, Yuen MMS, Jancsik S, Madilao LL, Page JE, Bohlmann J (2020) Terpene Synthases and Terpene Variation in Cannabis sativa. Plant Physiol 184: 130–147** Google Scholar: <u>Author Only Title Only Author and Title</u>

**Braich S, Baillie RC, Jewell LS, Spangenberg GC, Cogan NOI (2019) Generation of a Comprehensive Transcriptome Atlas and Transcriptome Dynamics in Medicinal Cannabis. Sci Rep 9: 16583** Google Scholar: <u>Author Only Title Only Author and Title</u>

**Daniel B, Konrad B, Toplak M, Lahham M, Messenlehner J, Winkler A, Macheroux P (2017) The family of berberine bridge enzyme-like enzymes: A treasure-trove of oxidative reactions. Arch Biochem Biophys 632: 88–103** Google Scholar: <u>Author Only Title Only Author and Title</u>

**Frazee AC, Pertea G, Jaffe AE, Langmead B, Salzberg SL, Leek JT (2015) Ballgown bridges the gap between transcriptome assembly and expression analysis. Nat Biotechnol 33: 243–246** Google Scholar: <u>Author Only Title Only Author and Title</u>

**Fulvio F, Paris R, Montanari M, Citti C, Cilento V, Bassolino L, Moschella A, Alberti I, Pecchioni N, Cannazza G, et al (2021) Analysis of Sequence Variability and Transcriptional Profile of Cannabinoid synthase Genes in Cannabis sativa L. Chemotypes with a Focus on Cannabichromenic acid synthase. Plants 10: 1857** Google Scholar: <u>Author Only Title Only Author and Title</u>

**Grant CE, Bailey TL, Noble WS (2011) FIMO: scanning for occurrences of a given motif. Bioinformatics 27: 1017–1018** Google Scholar: <u>Author Only Title Only Author and Title</u>

**Grassa CJ, Weiblen GD, Wenger JP, Dabney C, Poplawski SG, Timothy Motley S, Michael TP, Schwartz CJ (2021) A new Cannabis genome assembly associates elevated cannabidiol (CBD) with hemp introgressed into marijuana. New Phytologist 230: 1665–1679** Google Scholar: <u>Author Only Title Only Author and Title</u>

**Gülck T, Møller BL (2020) Phytocannabinoids: Origins and Biosynthesis. Trends Plant Sci 25: 985–1004** Google Scholar: <u>Author Only Title Only Author and Title</u>

**Hanuš LO, Meyer SM, Muñoz E, Taglialatela-Scafati O, Appendino G (2016) Phytocannabinoids: a unified critical inventory. Nat Prod Rep 33: 1357–1392** Google Scholar: <u>Author Only Title Only Author and Title</u>

**Henikoff S, Henikoff JG (1992) Amino acid substitution matrices from protein blocks. Proceedings of the National Academy of Sciences 89: 10915–10919** Google Scholar: <u>Author Only Title Only Author and Title</u>

**Hurgobin B, Tamiru-Oli M, Welling MT, Doblin MS, Bacic A, Whelan J, Lewsey MG (2021) Recent advances in Cannabis sativa genomics research. New Phytologist 230: 73–89** Google Scholar: <u>Author Only Title Only Author and Title</u>

**Kojoma M, Seki H, Yoshida S, Muranaka T (2006) DNA polymorphisms in the tetrahydrocannabinolic acid (THCA) synthase gene in “drug-type” and “fiber-type” Cannabis sativa L. Forensic Sci Int 159: 132–140** Google Scholar: <u>Author Only Title Only Author and Title</u>

**Larkin MA, Blackshields G, Brown NP, Chenna R, McGettigan PA, McWilliam H, Valentin F, Wallace IM, Wilm A, Lopez R, et al (2007) Clustal W and Clustal X version 2.0. Bioinformatics 23: 2947–2948** Google Scholar: <u>Author Only Title Only Author and Title</u>

**Laverty KU, Stout JM, Sullivan MJ, Shah H, Gill N, Holbrook L, Deikus G, Sebra R, Hughes TR, Page JE, et al (2019) A physical and genetic map of Cannabis sativa identifies extensive rearrangements at the THC/CBD acid synthase loci. Genome Res 29: 146–156** Google Scholar: <u>Author Only Title Only Author and Title</u>

**Lazarini-Lopes W, do Val-da Silva RA, da Silva-Júnior RMP, Leite JP, Garcia-Cairasco N (2020) The anticonvulsant effects of cannabidiol in experimental models of epileptic seizures: From behavior and mechanisms to clinical insights. Neurosci Biobehav Rev 111: 166–182** Google Scholar: <u>Author Only Title Only Author and Title</u>

**Li H, Handsaker B, Wysoker A, Fennell T, Ruan J, Homer N, Marth G, Abecasis G, Durbin R (2009) The Sequence Alignment/Map format and SAMtools. Bioinformatics 25: 2078–2079** Google Scholar: <u>Author Only Title Only Author and Title</u>

**Li MZ, Elledge SJ (2012) SLIC: A method for sequence- and ligation-independent cloning. Methods in Molecular Biology 852: 51– 59** Google Scholar: <u>Author Only Title Only Author and Title</u>

**Liu Y, Zhu P, Cai S, Haughn G, Page JE (2021) Three novel transcription factors involved in cannabinoid biosynthesis in**

**Cannabis sativa L. Plant Mol Biol 106: 49–65** Google Scholar: <u>Author Only Title Only Author and Title</u>

**Lynch RC, Vergara D, Tittes S, White K, Schwartz CJ, Gibbs MJ, Ruthenburg TC, deCesare K, Land DP, Kane NC (2016) Genomic and Chemical Diversity in Cannabis. CRC Crit Rev Plant Sci 35: 349–363** Google Scholar: <u>Author Only Title Only Author and Title</u>

**McLeay RC, Bailey TL (2010) Motif Enrichment Analysis: a unified framework and an evaluation on ChIP data. BMC Bioinformatics 11: 165** Google Scholar: <u>Author Only Title Only Author and Title</u>

**McPartland JM, Hegman W, Long T (2019) Cannabis in Asia: its center of origin and early cultivation, based on a synthesis of subfossil pollen and archaeobotanical studies. Veg Hist Archaeobot 28: 691–702** Google Scholar: <u>Author Only Title Only Author and Title</u>

**de Meijer EPM, Bagatta M, Carboni A, Crucitti P, Moliterni VMC, Ranalli P, Mandolino G (2003) The Inheritance of Chemical Phenotype in Cannabis sativa L. Genetics 163: 335–346** Google Scholar: <u>Author Only Title Only Author and Title</u>

**Michael Felberbaum (2018) https://www.fda.gov/news-events/press-announcements/fda-approves-first-drug-comprised-active-ingredient-derived-marijuana-treat-rare-severe-forms. https://www.fda.gov/news-events/press-announcements/fda-approves-first-drug-comprised-active-ingredient-derived-marijuana-treat-rare-severe-forms,** Google Scholar: <u>Author Only Title Only Author and Title</u>

**Morimoto S, Komatsu K, Taura F, Shoyama Y (1998) Purification and characterization of cannabichromenic acid synthase from Cannabis sativa. Phytochemistry 49: 1525–1529** Google Scholar: <u>Author Only Title Only Author and Title</u>

**Napiroon T, Tanruean K, Poolprasert P, Bacher M, Balslev H, Poopath M, Santimaleeworagun W (2021) Cannabinoids from inflorescences fractions of Trema orientalis (L.) Blume (Cannabaceae) against human pathogenic bacteria. PeerJ 9: e11446** Google Scholar: <u>Author Only Title Only Author and Title</u>

**Onofri C, de Meijer EPM, Mandolino G (2015) Sequence heterogeneity of cannabidiolic- and tetrahydrocannabinolic acid-synthase in Cannabis sativa L. and its relationship with chemical phenotype. Phytochemistry 116: 57–68** Google Scholar: <u>Author Only Title Only Author and Title</u>

**Pagès H, Aboyoun P, Gentleman R, DebRoy S (2020) Biostrings: Efficient manipulation of biological strings.**

**Pauli CS, Conroy M, vanden Heuvel BD, Park S-H (2020) Cannabidiol Drugs Clinical Trial Outcomes and Adverse Effects. Front**

**Pharmacol. doi: 10.3389/fphar.2020.00063** Google Scholar: <u>Author Only Title Only Author and Title</u>

**Pertea M, Kim D, Pertea GM, Leek JT, Salzberg SL (2016) Transcript-level expression analysis of RNA-seq experiments with**

**HISAT, StringTie and Ballgown. Nat Protoc 11: 1650–1667** Google Scholar: <u>Author Only Title Only Author and Title</u>

**Reid G (1991) Molecular cloning: A laboratory manual, 2nd edn by J. Sambrook, E. F. Fritsch and T. Maniatis, Cold Spring Harbor Laboratory Press, 1989. $115.00 (3 vols; 1659 pages) ISBN 0 87969 309 6. doi: 10.1016/0167-7799(91)90068-S** Google Scholar: <u>Author Only Title Only Author and Title</u>

**Ren G, Zhang X, Li Y, Ridout K, Serrano-Serrano ML, Yang Y, Liu A, Ravikanth G, Nawaz MA, Mumtaz AS, et al (2021) Large-scale whole-genome resequencing unravels the domestication history of Cannabis sativa. Sci Adv. doi: 10.1126/sciadv.abg2286** Google Scholar: <u>Author Only Title Only Author and Title</u>

**Shoyama Y, Tamada T, Kurihara K, Takeuchi A, Taura F, Arai S, Blaber M, Shoyama Y, Morimoto S, Kuroki R (2012) Structure and Function of Δ1-Tetrahydrocannabinolic Acid (THCA) Synthase, the Enzyme Controlling the Psychoactivity of Cannabis sativa. J Mol Biol 423: 96–105** Google Scholar: <u>Author Only Title Only Author and Title</u>

**Sirikantaramas S, Morimoto S, Shoyama Y, Ishikawa Y, Wada Y, Shoyama Y, Taura F (2004) The Gene Controlling Marijuana**

**Psychoactivity. Journal of Biological Chemistry 279: 39767–39774** Google Scholar: <u>Author Only Title Only Author and Title</u>

**Slater G, Birney E (2005) Automated generation of heuristics for biological sequence comparison. BMC Bioinformatics 6: 31** Google Scholar: <u>Author Only Title Only Author and Title</u>

**Stack GM, Toth Ja, Carlson CH, Cala AR, Marrero-González MI, Wilk RL, Gentner DR, Crawford JL, Philippe G, Rose JKC, et al (2021) Season-long characterization of high-cannabinoid hemp (Cannabis sativa L.) reveals variation in cannabinoid accumulation, flowering time, and disease resistance. GCB Bioenergy 13: 546–561** Google Scholar: <u>Author Only Title Only Author and Title</u>

**Taura F, Sirikantaramas S, Shoyama Y, Yoshikai K, Shoyama Y, Morimoto S (2007) Cannabidiolic-acid synthase, the chemotype-determining enzyme in the fiber-type Cannabis sativa. FEBS Lett 581: 2929–2934** Google Scholar: <u>Author Only Title Only Author and Title</u>

**Thomas F, Kayser O (2022) Natural deep eutectic solvents enhance cannabinoid biotransformation. Biochem Eng J. doi: 10.1016/j.bej.2022.108380** Google Scholar: <u>Author Only Title Only Author and Title</u>

**Toth JA, Stack GM, Cala AR, Carlson CH, Wilk RL, Crawford JL, Viands DR, Philippe G, Smart CD, Rose JKC, et al (2020) Development and validation of genetic markers for sex and cannabinoid chemotype in Cannabis sativa L. GCB Bioenergy 12: 213–222** Google Scholar: <u>Author Only Title Only Author and Title</u>

**van Velzen R, Schranz ME (2021) Origin and Evolution of the Cannabinoid Oxidocyclase Gene Family. Genome Biol Evol 13: 2020.12.18.423406** Google Scholar: <u>Author Only Title Only Author and Title</u>

**Vergara D, Bidwell LC, Gaudino R, Torres A, Du G, Ruthenburg TC, deCesare K, Land DP, Hutchison KE, Kane NC (2017) Compromised External Validity: Federally Produced Cannabis Does Not Reflect Legal Markets. Sci Rep 7: 46528** Google Scholar: <u>Author Only Title Only Author and Title</u>

**Weiblen GD, Wenger JP, Craft KJ, ElSohly MA, Mehmedic Z, Treiber EL, Marks MD (2015) Gene duplication and divergence affecting drug content in Cannabis sativa. New Phytologist 208: 1241–1250** Google Scholar: <u>Author Only Title Only Author and Title</u>

**Wickham H (2016) ggplot2: Elegant Graphics for Data Analysis.**

**Zager JJ, Lange I, Srividya N, Smith A, Lange BM (2019) Gene Networks Underlying Cannabinoid and Terpenoid Accumulation in Cannabis. Plant Physiol 180: 1877–1897** Google Scholar: <u>Author Only Title Only Author and Title</u>

**Zirpel B, Kayser O, Stehle F (2018) Elucidation of structure-function relationship of THCA and CBDA synthase from Cannabis sativa L. J Biotechnol 284: 17–26** Google Scholar: <u>Author Only Title Only Author and Title</u>

